# Oligodendrocyte lineage cells driven by neuronal activity in selected brain regions are required for episodic memory formation

**DOI:** 10.1101/2021.12.10.472135

**Authors:** Luendreo P. Barboza, Benjamin Bessières, Omina Nazarzoda, Cristina M. Alberini

**Affiliations:** Center for Neural Science, New York University, New York; Neuroscience and Physiology, Neuroscience Institute, New York University Langone Medical Center, New York; School of Medicine, University of Virginia, Virginia; School of Natural and Behavioral Sciences, CUNY Brooklyn College, New York

**Keywords:** episodic memory, oligodendrocyte lineage cells, hippocampus, anterior cingulate cortex, rat, mouse

## Abstract

The formation of long-term episodic memories requires the activation of molecular mechanisms in several regions of the medial temporal lobe, including the hippocampus and anterior cingulate cortex (ACC). The extent to which these regions engage distinct mechanisms and cell types to support memory formation is not well understood. Recent studies reported that oligodendrogenesis is essential for learning and long-term memory; however, whether oligodendrocyte lineage cells are required only in selected brain regions is still unclear. Also still unknown are the temporal kinetics of oligodendrocyte lineage cells involvement in memory processes and whether these cells are engaged in response to neuronal activity. Here we show that in rats and mice, episodic learning rapidly increases the oligodendrogenesis and myelin biogenesis transcripts *Olig2*, *Myrf*, *Mbp*, and *Plp1* as well as oligodendrocyte precursor cells (OPC) proliferation and differentiation in the ACC, but not in the dorsal hippocampus (dHC). Region-specific knockdown or knockout of *Myrf*, a regulator of oligodendrocyte differentiation, revealed that cells of the oligodendrocyte lineage are required for memory formation in the ACC but not the dHC. Chemogenetic neuronal silencing in the ACC showed that neuronal activity is critical for learning-induced OPC proliferation. Hence, activity-driven oligodendrocyte lineage cells in the ACC, but not dHC, are critical for the formation of episodic memories.

**Impact statement:** Oligodendrocyte lineage cells are required in the anterior cingulate cortex but not in the hippocampus for long-term memory formation.

## Introduction

Long-term memories are initially fragile but become resilient to disruption through consolidation, a temporally graded process that requires cascades of molecular changes in select brain regions. Episodic memories become consolidated by rapidly recruiting molecular pathways in several brain regions, including the hippocampus, medial prefrontal cortex (mPFC), and anterior cingulate cortex (ACC) (Frankland & Bontempi, 2005; Kandel et al., 2014; Squire et al., 2015). In rodents, the molecular cascades underlying memory consolidation in the hippocampus are required for days (Bambah-Mukku et al. 2014), whereas in cortical regions they continue to play a critical role for weeks (Chen et al., 2020; Heyward & Sweatt, 2015), suggesting that a differential engagement of biological regulations in different regions support memory consolidation and storage. This distinctive regional regulation is likely the consequence of their different cellular and biological composition, which is also distinctively recruited in response to learning (Chen et al., 2020; Katzman et al., 2021; Saunders et al., 2018).

Although research in the field of learning and memory has thus far mostly focused on neuronal mechanisms and circuitry, in the last decade it has become clear that long-term memory formation requires the contribution of multiple cell types, including astrocytes (Adamsky & Goshen, 2018; Gerlai et al., 1995; Suzuki et al., 2011), microglia (Morris et al., 2013; Yirmiya & Goshen, 2011), and oligodendrocytes (McKenzie et al., 2014; Xin & Chan, 2020). Recent reports showed that oligodendrogenesis and *de novo* myelination play critical roles in the formation of several types of memory, including motor, spatial, and episodic (McKenzie et al., 2014; Pan et al., 2020; Steadman et al., 2020; Wang et al., 2020) as well as in sensory enrichment (Hughes et al., 2018). These studies examined the role of oligodendrocytes by assessing brain-wide oligodendrogenesis, however, several questions remain to be addressed: First, are distinct brain regions differentially engaging oligodendrocyte lineage cells and mechanisms in learning and memory? Second, what is the temporal progression of oligodendrocyte lineage cells requirement in the various processes of learning and memory? And, finally, do oligodendrocyte changes require neuronal activation?

Steadman et al. (2020) reported that the acquisition of spatial memory in mice is accompanied by an increase in oligodendrocyte precursor cell (OPCs) proliferation and/or differentiation mechanisms in the ACC, mPFC, and corpus callosum/cingulum (CC/Cg), but not in the hippocampus, suggesting that these regions may differentially involve oligodendrogenesis in memory formation. Yet, whether oligodendrocyte lineage cells are differentially engaged in different regions remains to be tested. In addition, using brain-wide conditional knockout of myelin regulatory factor (*Myrf*, a transcription factor required for oligodendrogenesis) in the oligodendrocyte precursors neuron-glial antigen 2 (NG2)-positive cells, the same authors provided evidence that oligodendrogenesis is required for long-term memory formation and learning-induced ripple-spindle coupling between the hippocampus and ACC, a cross-region synchronization believed to contribute to memory consolidation. Their temporal assessment of the effects of the conditional *Myrf* knockout on memory suggested that disrupting oligodendrogenesis during learning or the initial phase of memory consolidation disrupts both recent (tested 1 day later) and remote spatial memories (tested 28 days later) (Steadman et al., 2020), whereas *Myrf* knockout 25 days after training had no effect on memory retention. These results lead to the conclusion that spatial learning and/or consolidation, but not remote memory storage, requires oligodendrogenesis. Pan et al. (2020) obtained a different result using another hippocampus-dependent task in mice, contextual fear conditioning. These authors found that *Myrf* knockout in the NG2-positive cells prior to learning impairs remote memory (tested 30 days after training) but not recent memory (tested 1 day after training). Hence, while the effect of oligodendrogenesis on recent hippocampus-dependent memories is still under debate, both studies concluded that experience-dependent oligodendrogenesis is required for long-term memory formation. Notably, the contribution of oligodendrocyte mechanisms during acquisition of information, i.e., learning, remains to be defined. Finally, while previous work showed that neuronal activity promotes oligodendrogenesis and adaptive myelination (Baraban et al., 2016; Gibson et al., 2014), whether learning-induced oligodendrocyte lineage cell functions requires neuronal activity remains to be established.

To address these questions, we employed the episodic memory paradigm inhibitory avoidance (IA), in rats and mice, in order to test the effects across species and with different techniques. We found that OPC proliferation and oligodendrocyte differentiation in the ACC, but not dorsal hippocampus (dHC), is rapidly induced following learning and that oligodendrocyte lineage cells are required for memory consolidation, whereas they are dispensable for acquisition and storage of the memory. We also observed that learning-induced OPC proliferation in the ACC is dependent upon neuronal activity.

## Results

### Episodic learning rapidly increases oligodendrocyte-specific mRNAs and proteins in the ACC but not dHC of rats

To assess whether episodic learning is accompanied by oligodendrocyte-specific changes in the dHC and ACC of rats, we employed IA, a learning paradigm that results in long-term memory formation after a single-context-footshock association (Gold, 1986). We performed a time-course analysis of mRNAs typically expressed during oligodendrocyte differentiation and myelin biogenesis using reverse transcription–quantitative polymerase chain reaction (RT–qPCR) on ACC and dHC RNA extracts collected one hour, one day, and seven days following training (Fig. 1A). We analyzed the expression of *Myrf*, oligodendrocyte transcription factor 2 (*Olig2*), ectonucleotide pyrophosphatase 6 (*Enpp6*), myelin basic protein (*Mbp*), proteolipid protein 1 (*Plp1*), and myelin associated glycoprotein (*Mag*). *Olig2* is required for terminal differentiation of OPCs and indirectly induces the transcription of *Myrf*, a regulator of oligodendrocyte differentiation and myelin biogenesis (Bujalka et al., 2013; Emery, 2013). MYRF protein binds to the promoter regions of myelin-associated genes and regulates the transcription of *Mbp*, *Plp1*, and *Mag* (Bujalka et al., 2013). Together, the proteins MAG, PLP1, and MBP ensure proper myelin biogenesis, wrapping, compaction, and function (Sherman & Brophy, 2005; Simons & Nave, 2015). ENPP6 is a choline phosphodiesterase involved in lipid metabolism and myelin biogenesis (Morita et al., 2016).

**Figure. 1.**
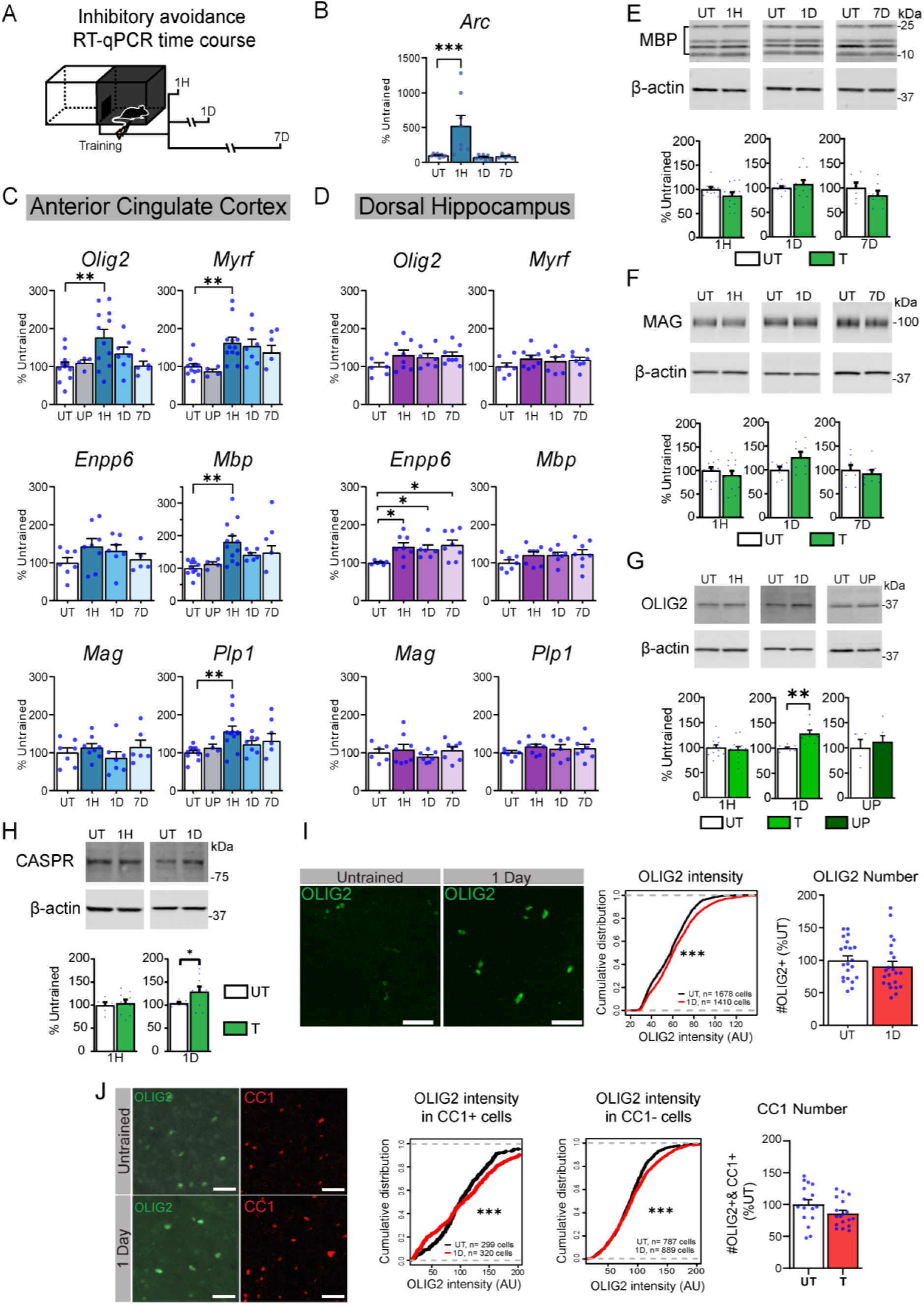
Learning rapidly induces oligodendrocyte-specific mRNAs and proteins. (A) Schematic representation showing the experimental design: rats underwent IA training and were euthanized at 1 hour (1H), 1 day (1D), or 7 days (7D) after training and assessed with RT-qPCR. (B) RT-qPCR of *arc* performed in ACC extracts from untrained (UT) and trained rats euthanized at the timepoints indicated in A. (C) ACC and (D) dHC RT-qPCR analyses of OPC and oligodendrocyte-specific genes *Olig2*, *Myrf*, *Enpp6*, *Mbp*, *Mag* and *Plp1*. Data are expressed as mean percentage ± s.e.m. of the untrained group (UT). Unpaired (UP) controls were added in groups where significant upregulation of oligodendrocyte genes were found(n = 4-12 per group; one-way ANOVA followed by Dunnett’s multiple comparison test; 2 independent experiments). (E-H) Examples and densitometric western blot analyses of MBP, MAG, CASPR, and OLIG2 obtained from ACC total extracts from trained rats euthanized at time points mentioned in A, compared to respective age-matched UT controls. UP controls were included at the one-day post training timepoint. Data presented as mean percentage ± s.e.m. of untrained rats (n = 4-12 rats per group; two-tailed t-test; full blot images can be found in Source Data file 2). (I) Examples of immunofluorescent staining of OLIG2 in the ACC of rats euthanized 1D after IA compared to UT control. Cumulative distribution of OLIG2 intensity measured from nuclei of ACC from UT and trained rats perfused 1D after training (n = 1678 and 1410 cell across four rats in UT and 1D groups respectively; two-tailed t-test; P<0.001). Mean values ± s.e.m. of the total number of OLIG2+ cells—data presented as positive cells per mm^2^. Each dot represents the quantification of one image taken from the ACC. 4-6 images were taken per rat per side on a total of 4 rats [UT (n = 23) and Trained1D (n=20)]. (J) Examples of immunofluorescent staining of OLIG2 and CC1 in the ACC of rats euthanized 1D after IA training compared to UT control. Mean values ± s.e.m. of the total number of OLIG2+ & CC1+ cells—data presented as positive cells as a percent of OLIG2+ cells. Each dot represents the quantification of one image taken from the ACC. 4-6 images were taken per rat per side on a total of 3 rats pr group. [UT (n = 16) and Trained1D (n=18)]. Cumulative distribution of OLIG2 intensity measured from nuclei of OLIG2+ & CC1+ cells from ACC from UT and trained rats perfused 1D after training (n = 299 and 320 cell across three rats in UT and 1D groups respectively; two-tailed t-test; P<0.001). Cumulative distribution of OLIG2 intensity measured from nuclei of OLIG2+ & CC1-cells from ACC from UT and trained rats perfused 1D after training (n = 787 and 889 cell across three rats in UT and 1D groups respectively; two-tailed t-test; P<0.001); two-tailed t-test; * indicates p<0.05, ** indicates p<0.01*** indicates p<0.001. For detailed statistical information, see Table 1-Source Data1.

We first confirmed that trained rats exhibited brain activation by assessing the expression of the immediate-early gene *Arc* (Bramham et al., 2010; Okuno et al., 2012; Shepherd & Bear, 2011), and observed that, as expected, its mRNA expression was significantly increased one hour after training and back to baseline levels at one and seven days after training (Fig. 1B). The results of the qPCRs showed a rapid and significant increase in the levels of *Olig2*, *Myrf*, *Mbp*, and *Plp1* mRNAs in the ACC (Fig. 1C) at 1 hour after training relative to untrained (UT) controls, which remained in the homecage, or unpaired control rats (UP), which underwent context and shock exposure in a temporally unpaired fashion (one hour apart, see methods) and were euthanized 1 hour after the shock. All mRNA transcripts returned to baseline levels at one day and remained at control levels seven days after training. Notably, no significant changes in oligodendrocyte-specific transcripts were detected in the UP group relative to the UT control group, indicating that the mRNA changes observed in trained rats were due to associative learning and not to novel context or shock presentation. Expression of *Olig2*, *Myrf*, *Plp1*, *Mag*, and *Mbp* over the same time course in the dHC did not change (Fig. 1D), although a significant increase in *Enpp6* was detected at 1 hour and persisted at 1 and 7 days after training.

To determine whether the changes in mRNA expression were detected also at the protein level, we used western blot analyses to measure the relative concentrations of OLIG2, MBP, and MAG and also added the axonal membrane protein CASPR (Einheber et al., 1997). We found that learning led to a significant increase in OLIG2 and CASPR at one day after training without changing the levels of MAG and MBP (Fig.1 E–H). We observed no differences in the UP rats euthanized one day later compared to the untrained control group, suggesting that the significant increase in OLIG2 was the result of associative learning. We then examined the increase in OLIG2 protein in the ACC by performing immunohistochemical staining (Fig. 1I). Analyses of OLIG2-positive nuclei per pre-defined area of the ACC revealed that training significantly increased the intensity but not the number of OLIG2+ nuclei at 24 hours after training. To better dissect whether training led to an increase in differentiating and mature oligodendrocytes we performed immunohistochemical double staining analyses of OLIG2 and adenomatous polyposis coli clone CC-1 (CC1), a marker employed to visualize newly differentiating as well as mature myelinating oligodendrocytes (Fletcher et al., 2021; McKenzie et al., 2014; Xiao et al., 2016). Representative pictures with increased OLIG2 intensity confirmed that the majority of CC1 positive cells express OLIG2 even though, in some cells, OLIG2 appeared at low levels (Supplementary Fig. 1A). We did not observe a change in total OLIG2+ & CC1+ cells in the ACC after learning; however, we detected an increase in OLIG2 intensity in both CC1+ and CC1-cell populations (Fig. 1J).

Collectively these data showed that IA learning leads to a rapid increase in OPC and oligodendrocyte-specific markers involved in different phases of oligodendrocyte cell lineage development in the ACC, but not dHC, suggesting an ACC-specific role of oligodendrocytes in memory processes.

### Training increases OPC proliferation and oligodendrocyte differentiation in the ACC but not dHC of mice

The induction of *Myrf* and OLIG2 in the rat ACC but not the dHC following training (Fig. 1) suggested that ACC undergoes a learning-dependent differential increase in OPC proliferation and oligodendrocyte differentiation. To test this hypothesis, we quantified the rate of newly: i) dividing OPCs, ii) differentiating OPCs and iii) maturing oligodendrocytes in the ACC and dHC of mice. To label newly dividing cells, we injected intraperitoneally (*i.p*.) the thymidine analog 5-ethynyl-2’-deoxyuridine (EdU) into mice one hour before IA training and euthanized the mice one day later (Fig. 2A), a timepoint at which a significant training-dependent increase in OLIG2 protein levels was found (Fig. 1G and I). To determine the number of proliferating OPCs, we counted the number of cells doubly stained with fluorescent EdU and platelet-derived growth factor receptor alpha (PDGFRa), a marker of OPCs (Rivers et al., 2008). To determine the rate of newly differentiating oligodendrocytes, we measured the number of cells labelled with three markers, EdU, OLIG2 (a marker that labels all stages of oligodendrocyte lineage including OPCs) and CC1. To measure changes in number of maturing oligodendrocytes, we quantified the number of cells triply stained with EdU, OLIG2, and Glutathione-S-transferase (GST)-Pi, as GST-Pi is a marker of mature oligodendrocytes (Tansey et al., 1991). All these analyses were normalized against the total number of DAPI-positive cells. We found a significant increase in the number of EdU+ cells, EdU+&PDGFRa+ cells (Fig. 2B) and EdU+&OLIG2+&CC1+ (Fig. 2C) following IA training. Notably, EdU colocalized also with cells that had a low intensity CC1 immunofluorescence, which may reflect an initial stage of OPC differentiation following training. Additional quantifications of EdU+&OLIG2+&CC1+ cell number relative to the total number of OLIG2+ cells indicated that the training-dependent increase in triple staining takes place in less than 8% of oligodendrocyte lineage cells (Supplementary Fig. 1B). Furthermore, no significant change in the number of EdU-&OLIG2+&CC1+ was found between groups (Supplementary Data Fig. 1C), suggesting that training-related changes are detected only in the EdU+ cell population, hence proliferating OPCs. Finally, as expected by the limited temporal window of 24 h, which is likely not sufficient for the maturation of new proliferating OPCs into mature oligodendrocytes, a negligible number of EdU+&OLIG2+&GST-Pi+ cells (Fig. 2D) was detected in either untrained or trained conditions.

**Figure 2.**
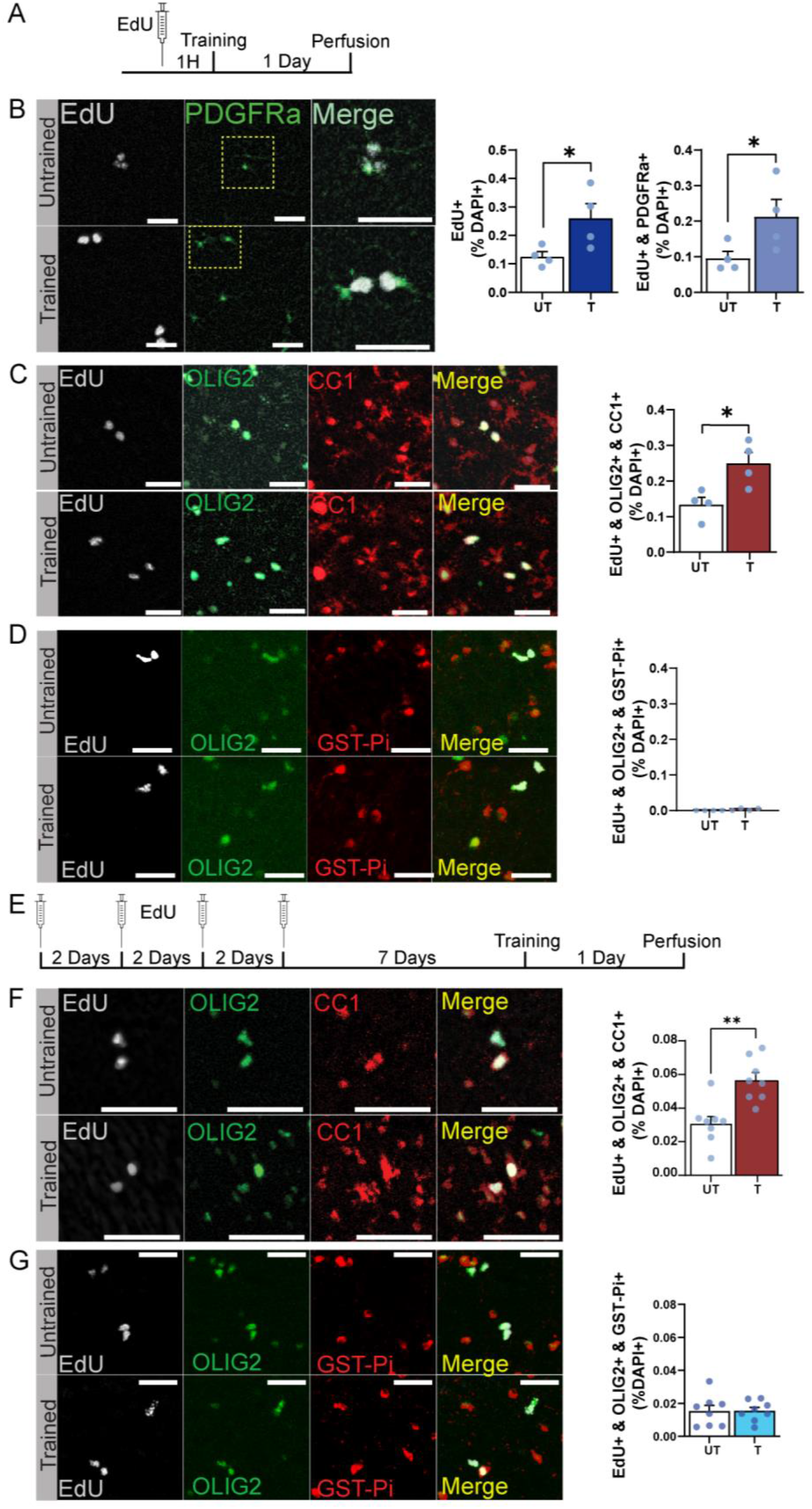
Training induces OPC proliferation and differentiation in the ACC. (A) Mice were injected with 5-ethynyl-2’-deoxyuridine (EdU) then one hour later they were trained in IA and finally perfused one day after training. Representative immunohistochemical staining of ACC sections and relative quantifications for (B) doubly stained EdU and PDGFRa cells; dotted box shown enlarged on the right side, triple staining of EdU, OLIG2 and CC1 and (D) triple staining of EdU, OLIG2 and GST-Pi. (E) Mice were injected with EdU once every other day for 4 injections terminating 7 days before training; they were then perfused one day after training. Representative immunohistochemical staining of ACC sections and relative quantifications for (F) EdU, OLIG2 and CC1, and (G) EdU, OLIG2 and GST-Pi. Each dot in the graphs represents the average value of three coronal section taken from each mouse. Data are expressed as mean percentage ± s.e.m. of positive cell number normalized against the total number of cells measured with DAPI+ staining (all scale bars: 40 μm; n = 4-8 mice per group, two-tailed t-test; *indicates P < 0.05, **indicates P < 0.01). For detailed statistical information, see supplementary table 2-Source Data1.

To extend the temporal window of detecting proliferating and differentiating OPCs, we labeled cells with EdU for a more extended period and also let them progress in their differentiation before testing their changes as a result of learning. Specifically, we injected EdU 4 times, once every other day, and then let the mice rest for 7 days before subjecting them to IA training. We then perfused the mice 1 day after training and performed immunohistochemistry (Fig. 2E). Similar to what found with EdU injected 1 hour before training, we detected a significant increase in EdU+&OLIG2+&CC1+ cells in the ACC of trained mice relative to untrained controls (Fig. 2F), but no significant change in EdU+ &OLIG2+ &GST-Pi+ cells (Fig. 2G). These results confirmed that there is a selective increase in newly differentiating OPCs (EdU+ & OLIG2+ & CC1+) but not maturation one day post-training.

Conversely, training did not change the number of EdU+, EdU+&PDGFRa+ or EdU+&OLIG2+&CC1+ cells neither in the entire dHC (Fig. 3A & B) nor in each hippocampal subregions DG, CA1, CA2, and CA3 (supplementary data Fig. 1D).

**Figure 3.**
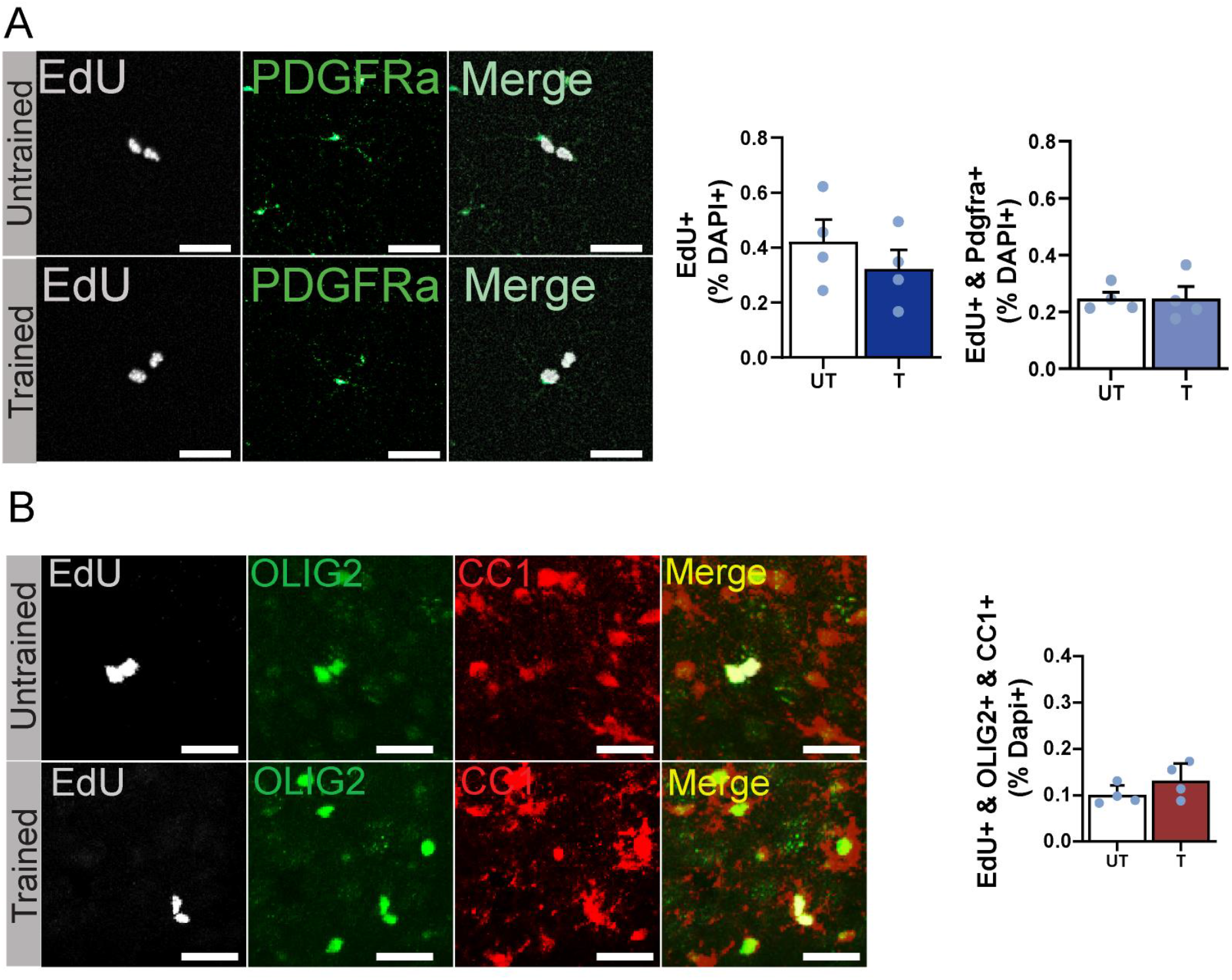
Training does not induce OPC proliferation and differentiation in the dHC. Mice were injected with 5-ethynyl-2’-deoxyuridine (EdU) and one hour later they were trained in IA and finally perfused one day after training. Representative immunohistochemical staining of dHP sections and relative quantifications for (A) doubly stained EdU and PDGFRa cells, (B) triple staining of EdU, OLIG2 and CC1. Each dot in the graphs represents the average value of three coronal section taken from each mouse. Data are expressed as mean percentage ± s.e.m. of positive cell number normalized against the total number of cells measured with DAPI+ staining (all scale bars: 40 μm; n = 4 mice per group, two-tailed t-test). For detailed statistical information, see supplementary table 3-Source Data1.

Collectively, these data indicated that IA training leads to a significant increase in proliferating OPCs and differentiating oligodendrocytes in the ACC but not dHC, which is detectable starting one day after training. However, newly proliferating OPCs did not undergo maturation in response to training during this temporal window.

### *Myrf* knockout disrupts memory formation

Next, we asked whether oligodendrogenesis is required for IA memory formation. We employed a conditional knockout mouse model in which *Myrf* is deleted in OPCs. Because MYRF is a transcription factor required for oligodendrocyte differentiation, its deletion in OPCs impairs oligodendrogenesis, and therefore new myelin formation, while leaving existing myelin unaffected (McKenzie et al., 2014). We tested the effect of *Myrf* knockout in OPCs on long-term memory using a double transgenic mouse line carrying the tamoxifen (TAM)-inducible CreER^T2^ expressed under the OPC-specific promoter *Pdgfra* and a floxed *Myrf* gene (*Pdgfra*-CreER^T2^ × *Myrf*^floxed/floxed^; hereafter, P-*Myrf*^floxed/floxed^). Injections of TAM in the P-*Myrf*^floxed/floxed^ mice temporally regulates the deletion of *Myrf* from OPCs. We used *Pdgfra*-CreER^T2^-*Myrf*^+/+^ (P-*Myrf*^+/+^) wild-type littermates as controls.

To confirm the effect of *Myrf* deletion on OPCs differentiation, TAM-treated P-*Myrf*^floxed/floxed^ and P-*Myrf*^+/+^ mice received an injection of EdU one hour before IA training and were perfused one day later. OPC differentiation was significantly inhibited in the ACC in P-*Myrf*^floxed/floxed^ mice, as demonstrated by the significant reduction in the number of cells positive for EdU, OLIG2, and CC1 in P-*Myrf*^floxed/floxed^ mice compared to P-*Myrf*^+/+^ littermate controls (Fig. 4A).

**Figure 4.**
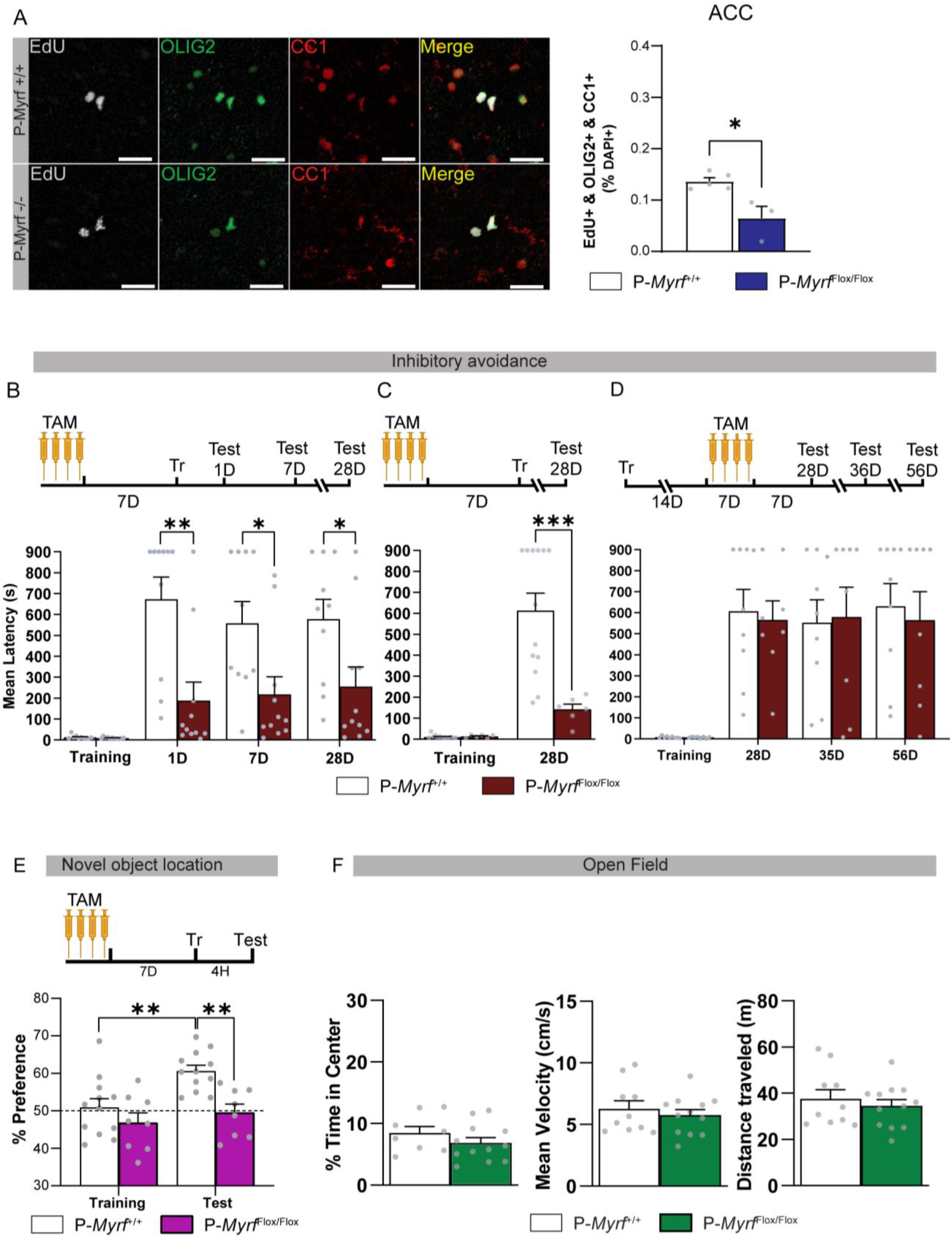
Conditional knockout of *Myrf* in OPCs impairs long-term memory. P-*Myrf*^+\+^ (*n* = 3) and P-*Myrf*^flox\flox^ (*n* = 5) littermates received one injection of tamoxifen (TAM) every other day for four times. Seven days after the last injection the mice underwent IA training. EdU was administered immediately before training and the mice were perfused one day after training. (A) Representative images and quantifications of ACC triple immunostaining (scale bar:40 μm) of EdU, OLIG2 and CC1. For each mouse, three coronal sections were quantified and averaged. In each coronal section the entire ACC was quantified bilaterally. Each dot represents the average of the three coronal section of each mouse. Data are presented as mean percentage ± s.e.m. of positive cell number relative to DAPI+ nuclei (scale bars: 40 μm;, two-tailed t-test). (B, C) P-*Myrf*^+\+^ and P-*Myrf*^flox\flox^ littermates were injected with TAM every other day for four injections terminating seven days before training. Mice were trained in IA and either tested at (B) 1, 7- and 28-days post-training (*n* = 10,11 per P-*Myrf*^+\+^ and P-*Myrf*^flox\flox^ per respectively) or (C) only at 28 days post-training *(n* = 13,6 per P-*Myrf*^+\+^ and P-*Myrf*^flox\flox^ groups respectively). (D) P-*Myrf*^+\+^ (*n* = 9) and P-*Myrf*^flox\flox^ (*n* = 8) littermates were trained and received tamoxifen injections starting 14 days after training and terminating seven days before testing, which occurred at 28D, 36D, and 56D post-training. Data are represented as mean latency ± s.e.m (In seconds, s). (two-way ANOVA followed by Bonferroni *post hoc* test). (E) P-*Myrf*^+\+^ (*n* = 12) and P-*Myrf*^flox\flox^ (*n* = 8) littermates were injected four times with tamoxifen once every other day. Seven days after the last injection the mice underwent novel object location training and were tested 4 hours later (two-way ANOVA followed by Bonferroni *post hoc* test). (F) Open field test expressed as mean ±s.e.m. of (i) percent time spent in the center of the arena, (ii) total distance, and (iii) mean velocity exploring the arena. (n = 9,12 mice per P-*Myrf*^+\+^ and P-*Myrf*^flox\flox^ groups respectively, two-tailed t-test; * indicates p<0.05, ** indicates p<0.01, *** indicates p<0.001). For detailed statistical information, see Table 4-Source Data1.

To test the effect of *Myrf* knockout on memory formation, TAM was administered to *Myrf*^floxed/floxed^ and P-*Myrf*^+/+^ mice seven days before IA training, and the mice were tested at 1, 7, and 28 days after training. P-*Myrf*^floxed/floxed^ mice exhibited significant memory reduction at all time points compared to P-*Myrf*^+/+^ controls (Fig. 4B). To exclude the potential effects of multiple testing, a second experiment was conducted in which P-*Myrf*^floxed/floxed^ and P-*Myrf*^+/+^ littermates were tested only at 28 days after training, and we again observed significant impairment in memory retention (Fig. 4C). We concluded that brain-wide oligodendrogenesis is required for the formation of recent and remote long-term memories.

To determine whether oligodendrogenesis contributes to the persistence or storage of memory, we administered TAM to P-*Myrf*^floxed/floxed^ and P-*Myrf*^+/+^ mice 14 days after training, when the consolidation process has significantly advanced (Bambah-Mukku et al., 2014; Squire et al., 2015). Memory retention was tested 14 days after knockout, corresponding to 28 days after training, as well as at 36 days and 56 days after training. No difference was detected between groups (Fig. 4D), indicating that oligodendrogenesis is not required for the persistence, retrieval, or storage of long-term memory.

Finally, to determine whether mechanisms involving oligodendrogenesis play a role in the formation of non-aversive episodic memories, P-*Myrf*^floxed/floxed^ and P-*Myrf*^+/+^ littermates were injected with TAM seven days before being trained in novel object location (nOL), a hippocampus-dependent learning paradigm (Mumby et al., 2002; Pezze et al., 2016; Weible et al., 2009) P-*Myrf*^floxed/floxed^ mice showed a significant nOL memory impairment compared to P-*Myrf*^+/+^ littermates (Fig. 4E) when tested four hours after training. Thus, oligodendrogenesis is also required for the formation of non-aversive hippocampus-dependent memories.

To exclude that the memory impairments we observed were due to other behavioral responses such as heightened anxiety-like responses or locomotor impairments, we tested P-*Myrf*^floxed/floxed^ and P-*Myrf*^+/+^ littermates in open field behavior. Time spent in the center of an open field arena and the distance and velocity traveled in the arena are putative measures of anxiety and locomotion abilities, respectively. No significant differences in anxiety-like and locomotor responses were detected; the time spent in the center and the distance and mean velocity traveled were similar between P-*Myrf*^floxed/floxed^ and P-*Myrf*^+/+^ littermates (Fig. 4F).

Collectively, these results indicate that oligodendrogenesis is required for the formation of long-term hippocampus-dependent memories.

### *Myrf* knockdown in the ACC but not the dHC of rats impairs memory consolidation but not learning

In order to investigate whether learning-dependent *Myrf* increase is differentially implicated in distinct brain regions and memory processes, we employed a *Myrf* knockdown strategy. We achieved region-specific and temporally restricted *Myrf* knockdown by using stereotactic injections to deliver an antisense oligodeoxynucleotide (ASO-ODN) specific against *Myrf* (Myrf-ASO), and, as a control, a related scrambled sequence (Myrf-SCR). We injected the ODNs bilaterally into the brain region of interest at various times before and after training.

The rapid and temporally limited effect of the ODN-mediated knockdown approach offers the opportunity to dissect, with highly defined temporal approaches, the dynamics of the requirement of specific mRNA translations, in addition to allowing the definition of anatomical requirements (Chen et al., 2011; Garcia-Osta et al., 2006; Taubenfeld et al., 2001). Hence, we used the Myrf-ASO approach to examine whether learning-induced expression of *Myrf* is required in the ACC and dHC for memory acquisition or consolidation.

To verify knockdown of *Myrf*, Myrf-ASO and Myrf-SCR were injected bilaterally 15 minutes before training, and *Myrf* mRNA levels were measured in the ACC one hour after training, when there is a significant learning-dependent increase in *Myrf* expression (Fig. 1C). Rats treated with Myrf-ASO had significantly lower *Myrf* mRNA levels compared to those treated with *Myrf*-SCR (Fig. 5A). Rats injected with Myrf-ASO exhibited no significant differences in MBP protein expression in the ACC one day after training, suggesting that Myrf-ASO treatment did not lead to demyelination (Fig. 5B).

**Figure 5.**
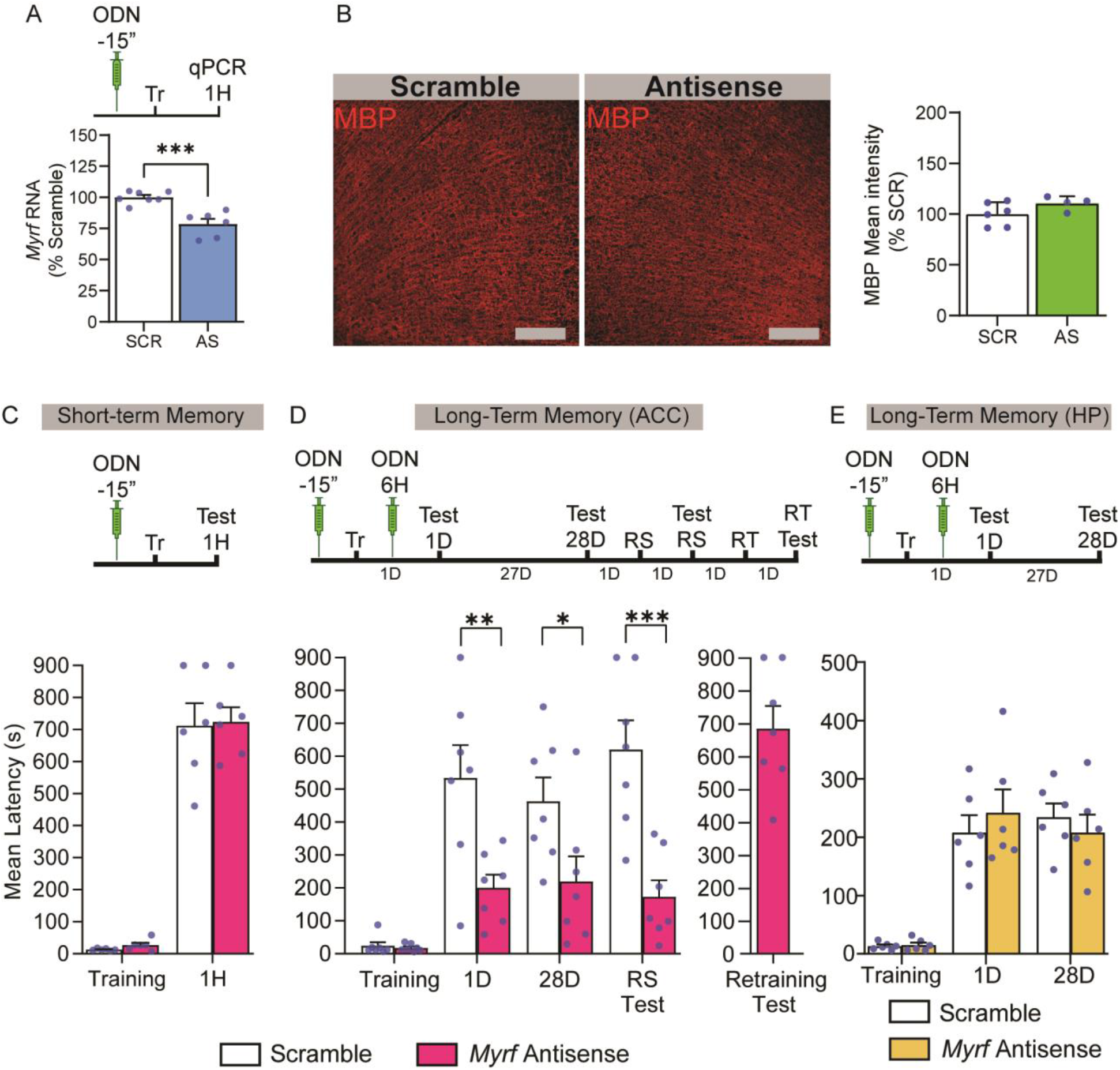
Antisense-mediated MYRF knockdown in the ACC impairs memory consolidation. (A) Rats were bilaterally injected in the ACC with either scrambled (SCR, *n* = 7) or antisense oligonucleotides against *Myrf* (ASO, *n* = 6) 15 minutes before training and euthanized one hour (1H) later for RT-qPCR analysis of *myrf* mRNA levels. (B) Immunohistochemistry representative images and quantification of MBP. Rats were bilaterally injected with either SCR (*n* = 6) or AS (*n* = 4) 15 minutes before and 6 hours after training and perfused one day after training for immunohistochemistry against MBP (scale bars: 160 μm). (C, D, E) Mean latency of rats in which ACC (C, D) and dHC (E) were bilaterally injected with either scramble sequences or *myrf* antisense oligonucleotides. (C) Injections were given 15 minutes before training and rats were tested at one hour post-training (*n* = 6 per group) or (D) injections were given 15 minutes before and 6 hours after training and rats were tested one day, and 28 days after training *(n* = 7 per group). Rats received a reminder shock (RS) after the last testing, followed by another retention test a day later (RS test). One day later rats underwent re-training (RT), and memory retention was tested a day later (RT test). (E) Injections were given 15 minutes before and 6 hours after training and rats were tested one day and 28 days after training *(n* = 6 per group). Data are presented as mean latency ± s.e.m. to enter the dark chamber (in seconds, s; two-way ANOVA followed by Bonferroni *post hoc* test; * indicates P < 0.05, ** indicates P < 0.01, *** indicates P < 0.001). For detailed statistical information, see table 5-source data1.

In order to test whether MYRF increase is required for learning, we bilaterally injected Myrf-ASO or Myrf-SCR into the ACC 15 minutes before training and tested the effect 1 hour after training. We detected no differences in memory between the two groups (Fig. 5C), indicating that MYRF is dispensable in the ACC for learning and short-term IA memory. To test whether MYRF is required for memory consolidation, bilateral injections of Myrf-ASO or Myrf-SCR were administered in the ACC 15 minutes before and six hours after training, then memory was tested one day after training. Rats injected with Myrf-ASO exhibited significant memory impairment one day after training compared to rats that had received Myrf-SCR injections (Fig. 5D), and the impairment persisted at 28 days after training (Fig. 5D). A reminder shock given one day after the remote memory test was unable to reinstate memory, indicating that the memory impairment was not due to a suppressed memory response but likely resulted from disrupted memory consolidation. Furthermore, re-training one day later of rats who had been injected with Myrf-ASO resulted in a long-lasting memory, thereby excluding the possibility that they had experienced memory loss because of ACC damage caused by surgery or injections.

By contrast, when Myrf-ASO was injected bilaterally into the dHC 15 minutes before and six hours after IA training, we observed no effect on memory retention; memories of the two treatment groups were similar at 1 day and 28 days after training (Fig. 5E). The lower level of retention in the dHC relative to the ACC with stereotactic injections is typically observed. Thus, we concluded that *Myrf* in the ACC is critical for the consolidation, but not the acquisition of IA, and is not required in the dHC.

### Oligodendrocytes in the mouse ACC are required for memory formation

Studies published thus far on the role of *Myrf* in memory formation have reported that brain-wide disruption of oligodendrogenesis impairs motor learning, spatial memory, and remote contextual fear memory in mice (McKenzie et al., 2014; Pan et al., 2020; Steadman et al., 2020). Our data in rats indicated that MYRF-dependent oligodendrocyte differentiation is essential for memory consolidation in the ACC but not the dHC. To investigate the region-specific roles of oligodendrocyte lineage cells in memory formation in the mouse brain, we bilaterally injected an adeno-associated viral vector expressing CreER^T2^ driven by the *Mbp* promoter (AAV-*Mbp*-CreER^T2^) in *Myrf*^+\+^ and *Myrf*^flox\flox^ mice to knock out *Myrf* selectively in oligodendrocytes of either ACC or dHC under the regulation of TAM. The choice of the *Mbp* promoter was twofold: first, our data showed an increase in *Mbp* mRNA following training; second, the *Mbp* promoter-based virus would have allowed us to test whether more mature stages of oligodendrocyte cell lineage functions, relative to OPC proliferation, are involved in memory formation. Notably, *Mbp* is expressed by premyelinating oligodendrocytes (Galiano et al., 2006) and found within a day of OPC differentiation (Dugas et al., 2006). Hence, we specifically tested whether MYRF-expressing differentiating and mature oligodendrocytes are required in the ACC and dHC for learning and/or memory formation. After two weeks to allow for viral expression, intraperitoneal injections of TAM were administered four times, once every other day, then mice underwent IA training (Fig. 6A). As the virus was not engineered with fluorophore sequences, we used diffusion of Chicago blue dye to confirm the injection diffusion area into the ACC (Fig. 6B). Compared to *Myrf*^+\+^ littermates, *Myrf*^flox\flox^ mice injected with AAV-*Mbp*-CreER^T2^ showed a significant decrease in the number of triply stained EdU+ & OLIG2+ & CC1+ cells indicating an effect of MYRF knockout on oligodendrocyte differentiation in the ACC (Fig. 6C) one day after training. There was no significant change in the number of EdU- & OLIG2+ & CC1+ cells between, *Myrf*^flox\flox^ and *Myrf*^+\+^ littermates (Supplementary Fig. 1E) suggesting that the effect of Myrf knockout at this timepoint is detected only in EdU-labeled cells. As EdU+ cells labeled with OLIG2 and CC1 constitute a small fraction of cells relative to their EdU-counterparts, we cannot exclude that changes also occur in the EdU-population but were undetectable due to insufficient sensitivity. Nevertheless, these results suggest that the AAV-*Mbp*-CreER^T2^ injection decreased differentiation mechanisms in *Mbp*-expressing oligodendrocytes validating the MYRF knockout.

**Figure 6.**
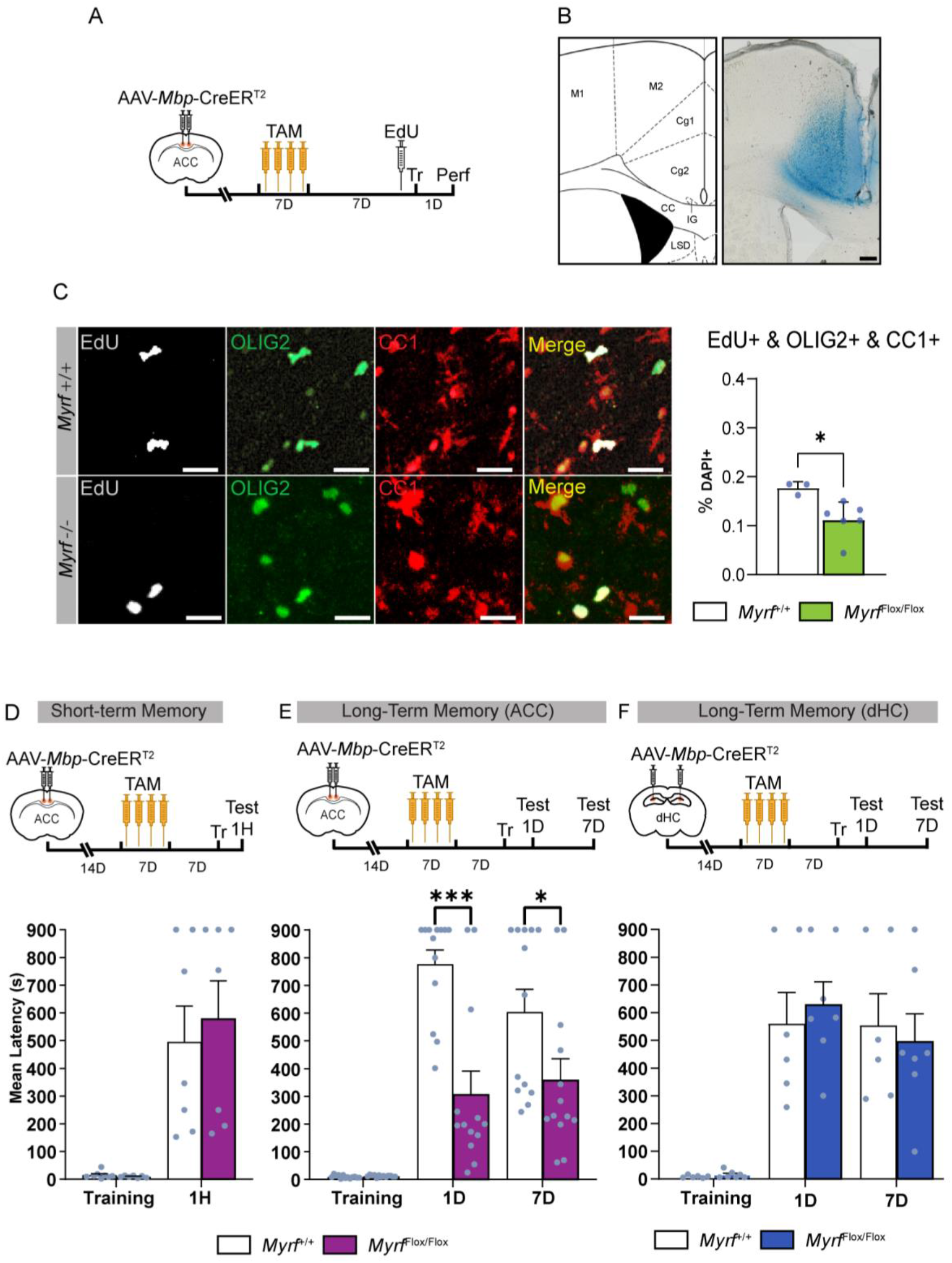
*Myrf* knockout in the mouse ACC impairs memory formation. (A) Experimental design: AAV-*Mbp*-CreER^T2^ was injected bilaterally into the ACC of *Myrf*^+\+^ (*n* = 3) and *Myrf*^flox\flox^ mice (*n* =5). Fourteen days following viral injection, mice were injected (*i.p.)* with tamoxifen (TAM) every other day for 4 times, terminating seven days before the training. Mice were then injected (*i.p.)* with EdU, immediately after IA training and perfused for immunohistochemistry one day after training. (B) Diffusion of injection shown by Chicago sky blue diffusion targeted mainly the ACC (right panel). Left panel image adapted from mouse brain atlas (scale bar: 200 μm). (C) Representative immunohistochemical staining (scale bar: 40 μm) and quantification of ACC triple immunostaining of EdU, OLIG2 and CC1. For each mouse, three coronal sections were quantified and averaged. In each coronal section the entire ACC was quantified bilaterally. Each dot represents the average of the three coronal sections of each mouse. Data are presented as mean percentage ± s.e.m. of positive cell number relative to DAPI+ nuclei in the ACC (two-tailed t-test). (D) AAV-*Mbp*-CreERT2 was injected bilaterally into the ACC of *Myrf*^+\+^, and *Myrf*^flox\flox^ mice. Fourteen days following viral injection, mice were received 4 injections of (TAM) (once every other day) and seven days later they underwent IA training and were tested at one hour (1H) after training to test short-term memory (*n* = 7 per group). (E) The mice underwent the same protocol described in D but were tested at one day (1D) and seven days (7D) after training to assess for long-term memory (*n* = 7,11 per *Myrf*^+\+^, and *Myrf*^flox\flox^ groups respectively). Data are represented as mean latency ± s.e.m. two-way ANOVA followed by Bonferroni *post hoc* test. (F) AAV-*Mbp*-CreER^T2^ was injected bilaterally into the dHC of *Myrf*^+\+^ (*n* = 6) and *Myrf*^flox\flox^ mice (*n* = 7). Fourteen days following viral injection, the mice received the 4 times TAM protocol and 7 days later were trained in IA. They were then tested at 1D and 7D after training. Data are represented as mean latency ±s.e.m. (in seconds, s; two-way ANOVA followed by Bonferroni *post hoc* test, * Indicates p<0.05, *** indicates p<0.001). For detailed statistical information, see table 6-source data1.

To test whether ACC-specific *Myrf* knockout is required for learning and short-term memory, another cohort of *Myrf*^+\+^ mice and *Myrf*^flox\flox^ littermates underwent similar viral injections and TAM protocol but were tested at one hour after IA training. No differences between groups were observed (Fig. 6D). To determine the role of oligodendrocytes in the ACC on memory processes, *Myrf*^+\+^ and *Myrf*^flox\flox^ littermates were treated with the same viral and TAM injection protocol as above but tested for memory retention at one day and seven days after training. Compared to *Myrf*^+\+^ littermates, *Myrf*^flox\flox^ mice showed a significant memory impairment at both time points after training (Fig. 6E). These results suggest that oligodendrocytes in the ACC are necessary for memory consolidation but dispensable for memory acquisition and short-term memory in mice, just as in rats.

To test whether oligodendrocytes is required for memory formation in the hippocampus, AAV-*Mbp*-CreER^T2^ was bilaterally injected into the dHC of *Myrf*^+\+^ and *Myrf*^flox\flox^ littermates using the protocol described above. No differences in memory retention were observed at one day or seven days post-training compared to control groups (Fig. 6F), leading us to conclude that oligodendrocytes are required in the ACC for memory consolidation but not for learning or short-term memory. By contrast, oligodendrocytes are dispensable in the dHC for the formation of hippocampus-dependent memories.

### DREADD-mediated neuronal inhibition impairs learning-induced OPC proliferation

Neuronal activity can drive OPC proliferation, oligodendrogenesis, and adaptive myelination (Gibson et al., 2014) however, it is not known whether neuronal activity is required to induce learning-dependent changes in oligodendrocyte lineage cells. To address this question, we employed the adeno-associated virus 8 (AAV8) expressing the Gi-coupled Designer Receptor Exclusively Activated by Designer Drugs (DREADD) hM4Di under the control of the human synapsin promoter to target expression to neurons (AAV-hSn-hM4D(Gi)-mCherry). We injected AAV-hSyn-hM4D(Gi)-mCherry bilaterally in the ACC and, after two weeks to allow for viral expression, we administered its DREADD ligand compound 21 (C21) *i.p*. one hour before IA training to transiently silence neuronal activity in the ACC (Fig. 7A) (Jendryka et al., 2019; Luo et al., 2021; Tran et al., 2020). In addition to hM4Di, the AAV-hSyn-hM4D(Gi)-mCherry viral construct expresses the fluorescent protein mCherry in neurons. Fluorescence was assessed by confocal microscopy two weeks after viral infection and found to be mostly confined to the ACC (Fig. 6B). The mice were tested one day after training. Treatment with C21 significantly impaired memory retention compared to vehicle injection (Fig. 7C), suggesting that neuronal activity in the ACC is required for memory formation.

**Figure 7.**
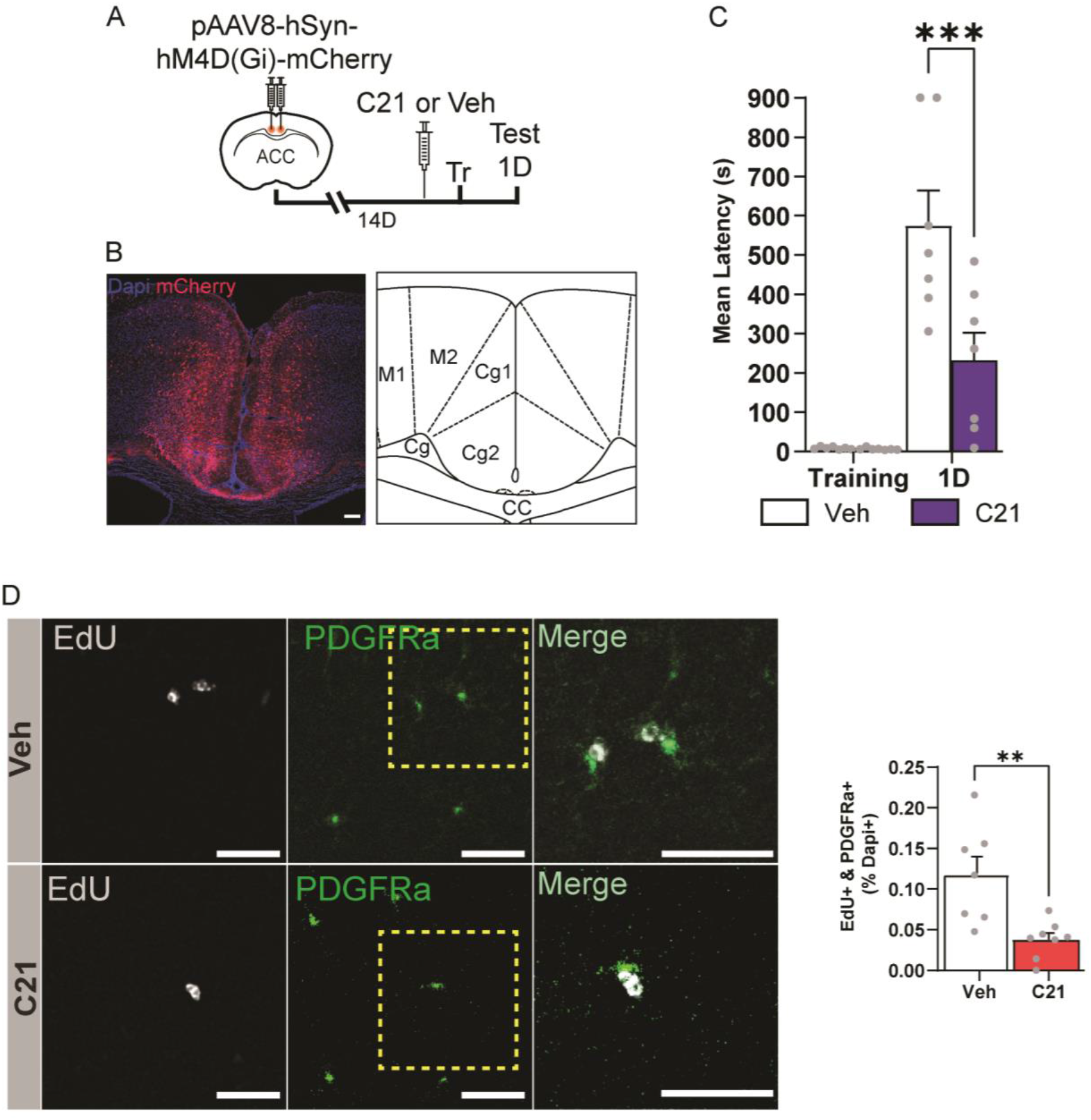
Neuronal activity inhibition in the ACC during learning impairs learning-dependent OPC proliferation and long-term memory formation. (A) Experimental schedule: AAV-hSyn-hM4D(Gi)-mCherry was injected bilaterally into the ACC of mice. Fourteen days following injection, the mice received an *i.p*. injection of C21 or Veh, and one hour later were trained in IA; the mice were tested one day after training. (B) Left panel: diffusion of AAV-hSyn-hM4D(Gi)-mCherry infection as revealed by mCherry fluorescence shows that it targeted the ACC (scale bar: 200 μm). Right panel: image adapted from mouse brain atlas. (C) Effect of C21 on IA memory: Mean latency ± s.e.m. of mice injected with either C21 or Vehicle (*n* = 7 per group) obtained at training and testing timepoints, expressed in seconds (s); two-way ANOVA followed by Bonferroni *post hoc* test) (D) Effect of C21 on OPC proliferation: AAV-hSyn-hM4D(Gi)-mCherry was injected targeting the ACC. Two weeks later mice were injected *i.p*. with 5-ethynyl-2’-deoxyuridine (EdU) and C21 or Veh, and one hour later underwent IA training; the mice were finally perfused 1 day after training. OPC proliferation in the ACC was assessed by double staining of EdU and PDGFRa. Left: representative image, scale bar: 40 μm; right: quantification of double staining. For each mouse, three coronal sections were quantified bilaterally and averaged. Each dot represents the average of the three coronal sections of each mouse. Data are expressed as mean percentage ± s.e.m. of positive cell number normalized against the total number of cells measured with DAPI+ staining; dotted box shown enlarged on the right side (n = 7-8 mice per group, two-tailed t-test; ** Indicates p<0.01, *** Indicates p<0.001). For detailed statistical information, see table 7-source data1.

To determine whether blocking neuronal activity in the ACC affected learning-dependent oligodendrocyte lineage cells we started by investigating OPC proliferation. Toward this end, AAV-hSyn-hM4D (Gi)-mCherry was bilaterally injected into the ACC and fourteen days later the mice were injected with either C21 or vehicle in combination with EdU one hour before receiving IA training. The mice were perfused one day after training at which point immunohistochemistry was used to quantify the number of cells that were positive for both EdU and PDGFRa. Trained mice injected with C21 had significantly fewer cells with EdU+&PDGFRA+ cells compared to mice injected with vehicle control, indicating that OPC proliferation was significantly impaired (Fig. 7D). Thus, we concluded that neuronal activity in the ACC is required for learning-induced changes in OPC proliferation.

## Discussion

This study showed that episodic learning, modeled by an IA paradigm in rats and mice, induces a rapid expression of the OPC and oligodendrocyte-specific mRNAs *Olig2*, *Myrf*, *Mbp*, and *Plp1* in the ACC but not in the dHC; however, an increase in *Enpp6* in the dHC was detected. The reason for this *Enpp6* increase is unclear; ENPP6 is a choline phosphodiesterase involved in lipid metabolism and myelin biogenesis (Morita et al., 2016); one possible explanation for its upregulation in the absence of changes in oligodendrogenesis markers is that, following learning, the hippocampus might regulate mechanisms of myelination and not oligodendrogenesis. In fact, whether existing myelin is remodeled after a learning experience is an open question.

Our western blot analyses confirmed that levels of OLIG2 significantly increased in the ACC following learning, supporting the idea that oligodendrogenesis is rapidly upregulated in this brain region in response to experience. Interestingly, MBP protein levels did not change, despite a significant increase in *Mbp* mRNA levels. This dichotomy might be due to the large pool of MBP in the brain, and it is possible that relatively small changes of MBP induced by a learning may not be easily detected with western blot. However, more sensitive techniques should be able to address this question. In line with the possibility of changes in MBP, both Pan et al. (2020) and Steadman et al. (2019) observed myelin changes in the cortex following episodic memory formation. Another oligodendrocyte-related protein, CASPR, which is an axonal membrane protein involved in myelin sheet growth, significantly increased after training in the ACC, confirming the idea that learning rapidly activates oligodendrocyte lineage cell-specific mechanisms in that region. The learning-induced changes at both mRNA and protein levels were found only with associative learning and were not detected in unpaired behavioral paradigms, which controlled for the separate experiences of context and footshock, indicating that oligodendrocyte-mediated mechanisms underlie associative memory processes. Notably, the types of oligodendrocyte lineage cell markers found to increase after learning suggest that several stages of oligodendrocyte differentiation, from oligodendrocyte precursor to mature myelinating oligodendrocytes, are regulated by experience.

Furthermore, our EdU labeling and immunohistochemistry experiments revealed that OPC proliferation (EdU+& PDGFRa+) and differentiation (EdU+&OLIG2+&CC1+) increased after learning in the ACC despite no change being detected in the total number of OLIG2+ cells. This lack of change in the number of OLIG2+ cells does not disprove the conclusion that learning induces oligodendrocyte differentiation, but rather suggests that the differentiation occurs either in only newly proliferating OPCs that undergo differentiation or in small cell populations that require more sensitive detection approaches. In fact, the EdU+ cell population accounts for a small number (< 0.5%) relative to the DAPI+ cell number of the ACC to which the quantifications were normalized. Even the analysis of the number of EdU+&OLIG2+&CC1+ cells relative to the total number of OLIG2+ cells in the ACC confirmed that the learning-dependent increase in OPC differentiation is detectable in the EdU+ but not in EdU-cell population. The EdU+ population is however less than 8% of OLIG2+ cells, again raising the question of whether learning-dependent changes occur either mostly in the proliferating OPCs, which rapidly proceed to differentiation, or also in other cell populations that however are too small to be detected with abundant markers of oligodendrogenesis stages.

Our results also extended previous findings on motor, spatial, and contextual fear memories by showing that brain-wide disruption of oligodendrogenesis impairs novel object location memories, strengthening the conclusion that oligodendrocyte lineage cells are required for long-term memory formation. Furthermore, by using multiple genetic and molecular approaches in rats and mice targeting specific brain regions of interest, we provided evidence that *Myrf* expression is required in the ACC but not the dHC, confirming the data across species. Thus, only certain brain regions in a given memory system recruit *Myrf* for memory consolidation.

Steadman et al. (2019) and Pan et al. (2020) recently reported that brain-wide *Myrf* knockout in OPCs prevents the formation of spatial and contextual memories. These studies showed that water maze and contextual fear conditioning learning in mice rapidly induce oligodendrocyte precursor cell (OPC) proliferation and differentiation into myelinating oligodendrocytes in cortical regions such as the ACC and medial prefrontal cortex (mPFC), but not the hippocampus. They suggested that myelin remodeling following training might be restricted to brain regions associated with long-term consolidation of hippocampus-dependent memories. However, because these studies utilized a brain-wide OPC knockout approach, they could not determine whether oligodendrogenesis in specific brain regions is required for memory formation. Identification of region- and circuitry-specific requirements for oligodendrogenesis and/or myelination in different types of learning and behavioral stimuli is important because it offers critical knowledge for better understanding the role of myelin in healthy brain functions as well as in diseases. Such knowledge should also expand our understanding of the circuitry that supports responses to learning.

Why oligodendrocyte lineage cells are required in the ACC but not the hippocampus is an open question; one possible explanation is that they may subserve long-term changes required for memory storage. In cortical regions including the ACC, but not in the hippocampus, episodic and spatial memories are stored for the very long term via systems consolidation, a process that requires time (Dudai et al., 2015). During systems consolidation the representation critical for memory expression/storage, which initially recruits both the hippocampus and cortical regions, undergoes a redistribution, and, while over time the hippocampus becomes dispensable, cortical regions such as ACC remain the site of long-term memory storage (Dudai et al., 2015; Frankland & Bontempi, 2005; Squire et al., 2015). Similarly, other types of memory such as motor memories are stored long-term through a consolidation process in cortical areas and precisely in motor cortices (Attwell et al., 2002; Krakauer & Shadmehr, 2006). More studies are needed to identify the region- and circuitry-specific recruitment of oligodendrogenesis and myelination underlying the consolidation of various memory systems and establish whether or not and how these mechanisms differentially serve different brain regions. For hippocampus-dependent memories, other cortical regions in addition to the ACC are likely to recruit oligodendrogenesis for memory formation and storage. For example, similarly to the ACC, the mPFC is involved in hippocampus-dependent long-term memory consolidation (Frankland & Bontempi, 2005), and the induction of oligodendrogenesis has been found in the mPFC after spatial and contextual fear learning (Pan et al., 2020; Steadman et al., 2020). Whether there are distinctive regulations of oligodendrogenesis between the ACC and the mPFC remains to be investigated.

In the agreement with findings that brain-wide oligodendrogenesis knockout impairs ripple-spindle coupling between the hippocampus and ACC (Steadman et al., 2020), the hippocampus may play an instructive role on the cortical oligodendrocyte lineage cell changes evoked by learning. Both Steadman et al. (2019) and Pan et al. (2020) suggested that experience-dependent myelination might promote the coupling of ensembles across regions to support the generation of a coordinated memory network because blocking *de novo* myelin formation throughout the brain, disrupted the activity and coordination in neural ensembles across the hippocampus and PFC networks.

Our results showing that learning increases OPC proliferation and oligodendrocyte differentiation targeted a limited temporal window after learning, i.e., 24 hours. Future studies extending the investigations to later time points should define if other stages of oligodendrocyte maturation including later stages of differentiation and myelination are also affected by learning. An extended time-course analysis should be able to also reveal whether learning-dependent myelination changes result from either the maturation of learning-induced OPC proliferation or take place in parallel as a result of activations of more mature populations of the oligodendrocyte lineage. Additionally, OPCs and non myelin-mediated functions of oligodendrocytes may also be involved in long-term memory formation via their abilities to regulate synaptic plasticity (Zemmar et al., 2018), neuronal excitability through extracellular potassium clearance (Larson et al., 2018), and trophic signals to neurons (Du & Dreyfus, 2002).

Our findings showing that changes of differentiation markers colocalized with markers of proliferation one day following IA learning are in line with previous studies showing that OPC differentiation is observed four hours after motor skill learning (Xiao et al., 2016) and required one day after water maze training (Steadman et al., 2020). Collectively these data suggest that the activation and recruitment of oligodendrocyte lineage cell mechanisms following learning are relatively rapid. Within a 24-hour temporal window we also detected a significant learning-dependent increase in *Mbp* mRNA in the ACC, which prompted us to test whether oligodendrogenesis and/or more mature oligodendrocytes played a critical role in memory. While the majority of *Mbp* expression is found in mature oligodendrocytes, *Mbp* is also expressed in differentiating/pre-myelinating oligodendrocytes, therefore, with the *Mbp*-driven knockout of MYRF, we could not fully dissect which stage of Mbp-expressing oligodendrocytes are critically recruited for memory formation. Nevertheless, we found that *Mbp*-regulated MYRF-expressing cells are required in the ACC for memory consolidation, further supporting the hypothesis that not only OPC proliferation but also oligodendrocyte differentiation in the ACC plays an important role in memory consolidation.

In the present study, we also dissected the requirement for *Myrf* in various memory processes. We found that training-induced *Myrf* expression in the ACC is necessary for the consolidation process but not for the initial acquisition of memory (learning) or remote storage. In fact, knocking down or knocking out *Myrf* before training did not affect short-term memory or acquisition, nor was there an effect on memory when *Myrf* was knocked down at a remote time point. However, disruption of oligodendrogenesis after training impaired long-term memory tested one day later, and the impairment persisted when the memory was tested at remote time points, such as four weeks after training. The lack of an effect on memory when oligodendrogenesis is disrupted weeks after training agrees with the results of Steadman et al. (2019), who showed that brain-wide OPC knockout of *Myrf* at 25 days after water maze training did not impair memory retention. From these results, we can conclude that MYRF-dependent changes in cortical regions are necessary for the rapid and initial phase of consolidation, but not for learning, retrieval, or memory storage.

Our results also shed light on the kinetics of oligodendrogenesis requirement in recent memory recall. Steadman et al. (2019) found that brain-wide *Myrf* conditional knockout in OPCs disrupts one-day-old spatial memory, and the disruption was still observed at a remote time point 28 days after training. By contrast, Pan et al. (2020) reported that *Myrf* knockout mice trained in contextual fear conditioning (CFC) had intact recent memory recall 1 day after training but impaired remote memories 28 days after training. We found that brain-wide and ACC-targeted knockout of *Myrf* in mice as well as ACC-specific ODN-mediated knockdown of MYRF in rats impaired recent memories tested one day after IA training. The impairments persisted in both rats and mice tested up to 28 days after training, leading us to conclude that MYRF-dependent oligodendrogenesis is rapidly upregulated and engaged following learning to selectively support a rapid phase of memory consolidation. It is possible that task-related differences in the kinetics of MYRF requirements exist, and that CFC has a slower cortical recruitment of oligodendrogenesis relative to water maze and IA tasks. It is also possible that *Myrf* knockout and knockdown under different conditions (ASO, *vs*. OPCs, *vs*. mature oligodendrocytes) and promoters may uncover distinct contributions of oligodendrocyte lineage cells to different memory phases. Knowing the role of oligodendrogenesis in specific memory processes and temporal phases of memory provides valuable information for the future development of temporally targeted treatments for cognitive symptoms of demyelinating diseases.

Finally, using a chemogenetic approach, we showed that the inhibition of neuronal activity in the ACC prevents learning-induced OPC proliferation. These findings extend previous data indicating that neuronal activity can drive changes in oligodendrocyte lineage cells (Baraban et al., 2016; Mount & Monje, 2017; Noori et al., 2020). We speculate that neurons that are activated during learning engage oligodendrocytes lineage cells at multiple stages to support the formation and storage of the memory long-term.

In sum, our data support the view that activity-regulated *Myrf*-dependent changes in oligodendrocyte lineage cells in select brain regions underlie long-term memory consolidation. We suggest that these changes support the stabilization process required to store information long-term.

## Materials and Methods

Key resources table:

| Reagent type<br>(species) or resource | Designation | Source or<br>reference | Identifiers | Concentration |
| --- | --- | --- | --- | --- |
| Strain, strain<br>background (R.<br>norvegicus, male) | CrI:LE Long-Evans | Charles River | Cat# 2308852,<br>RRID:RGD_23088<br>52 |  |
| Strain, strain<br>background (M.<br>Musculus, male and<br>female) | B6, B6J, B6/J | The Jackson<br>Laboratory | Stock No: 000664 |  |
| Strain, strain<br>background (M.<br>Musculus, male and<br>female) | B6.129S-Pdgrfra<sup>tm1.1(cre/ERT2)Blh/J</sup> | The Jackson<br>Laboratory | Stock No: 032770 |  |
| Strain, strain<br>background (M.<br>Musculus, male and<br>female) | B6;129-Myrf<sup>tm1Barr/J</sup> | The Jackson<br>Laboratory | Stock No: 010607 |  |
| Commercial assay or<br>kit | Qiagen RNeasy<br>Micro Kit | Qiagen | Cat# 74004 |  |
| Commercial assay or<br>kit | Qiagen QuantiTect<br>Rev. Transcription<br>Kit | Qiagen | Cat# 205311 |  |
| Commercial assay or<br>kit | Bio-Rad iQ SYBR<br>Green Supermix | Bio-Rad | Cat# 1708880 |  |
| Commercial assay or kit | Bradford protein assay kit | Bio-Rad | Cat# 5000201 |  |
| antibody | Anti-OLIG2 | Millipore company | Cat# MABN50 | WB: 1:1000;<br>IHC: 1:1000 |
| antibody | Anti-MBP | Millipore company | Cat# SKB3-05-675 | WB: 1:500;<br>IHC: 1:1000 |
| antibody | Anti-CASPR | Millipore company | Cat# MABN69 | WB: 1:2000 |
| antibody | Anti-MAG | Cell Signaling | Cat# 9043S | WB: 1:1000 |
| antibody | anti-OLIG2 | Millipore company | Cat# AB9610 | IHC: 1:1000 |
| antibody | anti-CC1 | Calbiochem | Cat# OP80 | IHC 1:500 |
| antibody | anti-PDGFRa | Cell signaling | Cat# 3164S | IHC: 1:1000 |
| antibody | Anti-GST-Pi | MBL international corporation | Cat# 312 | IHC: 1:1000 |
| Commercial assay or kit | Click-iT™ Plus EdU Cell Proliferation Kit | Thermo Fisher Scientific | Cat# C20086 |  |
| Chemical compound, drug | 5-ethynyl-2'-deoxyuridine | Biosynth | Cat# 087011802 | 80mg/kg |
| Chemical compound, drug | DAPI (4',6-Diamidino-2-Phenylindole, Dihydrochloride) | Life technologies | Cat# CD1306 | 1:10,000 |
| antibody | Alexa Fluor® 647 Goat Anti-Mouse IgG (H+L) | Invitrogen | Cat# A21236 | IF: 1:800 |
| antibody | Alexa Fluor® 488 Goat anti-Rabbit IgG (H+L) | Invitrogen | Cat# A11034 | IF: 1:800 |
| antibody | Alexa Fluor & reg;<br>568 goat anti-rabbit<br>IgG (H+L) | Invitrogen | Cat# A11036 | IF: 1:800 |
| Genetic reagent | pAAV-MBP-CreER<sup>T2</sup> | Vector Biolabs | Cat# VB1545 | AAV-PHP.B<br>capsid; titer:<br>10x13 GC/μl |
| Genetic reagent | AAV-hSyn-hM4D(Gi)-mCherry | Addgene | Cat# 50475-AAV8 | titer: 7×10<sup>12</sup><br>vg/mL; |
| Chemical compound,<br>drug | compound 21 | Hello Bio | Cat#: HB6124 | 1mg/kg |
| Chemical compound,<br>drug | Tamoxifen | Sigma-Aldrich | Cat# T5648 | 200mg/kg |
| Antibody | IRDye 680LT | LI-COR<br>Biosciences | Cat# 926-68050,<br>RRID:AB_278364<br>2 | WB (1:10000) |
| Antibody | IRDye 800CW (goat<br>anti-rabbit) | LI-COR<br>Biosciences | Cat# 926-32211,<br>RRID:AB_621843 | WB (1:10000) |
| Antibody | anti-β-actin | Santa Cruz<br>Biotechnology | <u>Cat# sc-47778</u><br><u>HRP,</u><br><u>RRID:AB_271418</u><br><u>9</u> | WB (1:20000) |
| Software, algorithm | Adobe Illustrator C6 | Adobe | https://www.adobe.com/products/illustrator.html |  |
| Software, algorithm | R |  | http://cran.r-project.org/ |  |
| Software, algorithm | Image Studio Lite | LI-COR<br>Biosciences | https://www.licor.com/bio/products/software/image_studio_lite/ |  |

### Rats

All animal procedures complied with the US National Institute of Health Guide for the Care and Use of Laboratory Animals and were approved by the New York University Animal Care Committees. Adult male Long–Evans (Charles Rivers, Wilmington, MA) rats weighing between 200 and 250 g were used. Animals were individually housed and maintained on a 12-h light/dark cycle. Experiments were performed during the light cycle. All rats were pair-housed and allowed ad libitum access to food and water and were handled for 3 minutes per day for 5 days before behavioral procedures. For all experiments, rats were randomly assigned to different groups. All protocols complied with the National Institutes of Health Guidelines for the Care and Use of Laboratory Animals and were approved by the Institutional Animal Care and Use Committee at New York University.

### Mice

Male and female *Pdgfra*-CreER^T2^-*Myrf* floxed (P-*Myrf*) mice were obtained by crossing *Pdgfra*-CreER^T2^ (The Jackson Laboratory, Bar Harbor, ME; B6.129S-*Pdgfra*^tm1.1(cre/ERT2)Blh^/J; Stock No: 032770), and *Myrf* floxed mice (The Jackson Laboratory, Bar Harbor, ME; B6;129-*Myrf*^tm1Barr^/J; Stock No: 010607). Breeding was designed to produce P-*Myrf*^+/+^, P-*Myrf*^flox/+^ P-*Myrf*^flox/flox^, *Myrf*^+/+^, *Myrf*^flox/+^ *Myrf*^flox/flox^ male and female littermates. Mice were bred in the animal facilities at New York University under a 12 h/12 h light/dark cycle (light on at 07.00 a.m.) with food and water ad libitum. After weaning, mice were group-housed (two to four per cage) in transparent plastic cages (31 × 17 × 14 cm) with free access to food and water. For inducing Cre-mediated knockout, all P-*Myrf* groups were administered 4 intraperitoneal (*i.p*.) injections of tamoxifen (TAM, Sigma-Aldrich St Louis, MO; Cat# T5648) dissolved in corn oil every other day at a dosage of 0.2g/kg of mouse. Mice were handled for 3 min per day for 5 days before behavioral procedures. For oligodendrogenesis experiments, c57/BL6 8–10-week males were used.

All mice were 8-10 weeks old at the start of behavioral assays. For all experiments, mice were randomly assigned to different groups. All protocols complied with the National Institutes of Health Guide for the Care and Use of Laboratory Animals and were approved by the Institutional Animal Care and Use Committee at New York University.

### Inhibitory Avoidance

The paradigm employed a chamber (Med Associates Inc., St. Albans, VT), which consisted of a rectangular Perspex box divided into a white light illuminated compartment and a dark black shock compartment (each 20.3 cm × 15.9 cm × 21.3 cm) separated by a door. The chamber was located in a sound-attenuated, red light illuminated room. During training and re-training sessions, the animal was placed in the lit compartment with its head facing away from the door. After 10 seconds (s) for rats and 30s for mice, the door automatically opened, allowing the animal access to the dark compartment. The door closed when the animal entered the dark compartment with all four limbs, and a foot shock (2 s, 0.9 mA in rats and 0.7mA in mice) was administered. The animal was removed from the dark compartment (10 s after the shock for rats and immediately after for mice) and returned to its home cage. Memory tests were performed at designated time points by placing the animal back in the lit compartment and measuring their latency to enter the dark compartment. For the mice, the door was initially open at the beginning of the test and remained open until the animals entered the dark compartment with all four limbs. For the rats, the door automatically opened 10 s after being placed in the lit compartment. Foot shocks were not administered during memory testing, and testing was terminated at 900s. Reminder foot shocks (R.S.), with identical duration and intensity to those used in training (i.e., 2 s, 0.9 mA), were administered in a novel, neutral chamber with transparent walls in a different experimental room. The animal was placed into the neutral chamber for 10s before receiving a single R.S. The animal was removed from the chamber immediately after the R.S. and returned to its home cage.

Control groups consisted of 1) untrained (U.T.) animals which were handled like the experimental but, instead of undergoing training, remained in their home cage, and 2) unpaired (UP) animals, which underwent the I.A. box exposure procedure without receiving a shock and, one hour later, given a foot shock immediately after being placed on the grid of the dark chamber and then immediately returned to the home cage.

### Real-time quantitative PCR (RT-qPCR)

The bilateral dorsal hippocampus or ACC was dissected into TRIzol (Invitrogen, Waltham, MA). Total RNA was extracted from each animal sample using RNeasy Micro Kit (Qiagen, Hilden, Germany cat# 74004) and reverse-transcribed using Qiagen QuantiTect Rev. Transcription Kit (Cat# 205311). RT-PCR was done using a BioRad CFX96 Touch Real-Time PCR machine. Twenty ng of the first-strand cDNA was subjected to PCR amplification using Bio-Rad iQ SYBR Green Supermix (Bio-Rad Laboratories, Hercules, CA; Cat# 1708880). Forty cycles of PCR amplification were performed: denaturing at check cycle 95°C for 15 s, annealing at 60°C for 30 s, and extension for 20 s at 72°C. Triplicates were performed for each cDNA sample. Delta-delta CT method was used to determine the relative quantification of gene expression in trained and unpaired groups compared to untrained animals. Primer sequences used: *Mbp* (Forward, 5′ GGCAAGGACTCACACACAAGAA 3′; Reverse, 5′ CTTGGGTCCTCTGCGACTTC 3′), *Plp1* (Forward, 5′ GCCAGAATGTATGGTGTTC 3′; Reverse, 5′ CAGCAATCATGAAGGTGAG 3′), *Myrf* (Forward, 5′ CCACATCAGCAGAACAAGTG 3′; Reverse, 5′ ACACGATAGGTGAGCATAGG 3′), *Mag* (Forward, 5′ CTGTGGTCGCCTTTG 3′; Reverse, 5′ GCTCTCAGTGACAATCC 3′), *Olig2* (Forward, 5′ CACGTCTTCCACCAAGAAAG 3′; Reverse, 5′ GTCCATGGCGATGTTGAG 3′), *Enpp6* (Forward, 5′ TGTGAGGTCCACCAGATG 3′; Reverse, 5′ CCCGATGTCGAATGACTTG 3′), *Erbb3* (Forward, 5′ CTGGCGTCTTTGGAACTG 3′; Reverse, 5′ GCAGACTGGAATCTTGATGG 3′), *Arc* (Forward, 5′ CCCTGCAGCCCAAGTTCAAG 3′; Reverse, 5′ GAAGGCTCAGCTGCCTGCTC 3′). *Erg1* (Forward, 5′ ACCTACCAGTCCCAACTCATC 3′; Reverse, 5′ GACTCAACAGGGCAAGCATAC 3′). *Cfos* (Forward, 5′ ATCCTTGGAGCCAGTCAAGA 3′; Reverse, 5′ ATGATGCCGGAAACAAGAAG 3′) and *Gapdh* (Forward, 5′GAACATCATCCCTGCATCCA 3′; Reverse 5′CCAGTGAGCTTCCCGTTCA 3′) was used as an internal control.

### Western Blot Analysis

Rats were euthanized, and their brains were quickly removed and snap-frozen with pre-chilled 2-methyl butane on dry ice. Dorsal hippocampal and ACC punches were obtained with a neuro punch (19 gauge; Fine Science Tools, Foster City, CA) from frozen brains mounted on a cryostat at −20°C and isolated the bilateral regions per animal (individual animal sample). Individual animal samples were homogenized in ice-cold RIPA buffer (50 mM Tris base, 150 mM NaCl, 0.1% SDS, 0.5% Na-deoxycholate, 1% NP-40) with protease and phosphatase inhibitors (0.5 mM PMSF, 2 mM DTT, 1 mM EGTA, 2 mM NaF, 1 μM microcystin, 1 mM benzamidine, 1 mM sodium orthovanadate, and Sigma-Aldrich protease and phosphatase inhibitor cocktails). Protein concentrations were determined using the Bio-Rad protein assay (Bio-Rad Laboratories, Hercules, CA, USA). Equal amounts of total protein extract per sample (20 μg) were resolved on denaturing SDS-PAGE gels and transferred to the Immobilon-FL Transfer membrane (Bio-Rad Laboratories, Hercules, CA, USA) by electroblotting. Membranes were dried, reactivated in methanol, washed with water, and then blocked in the Biorad blocking buffer for 2 h at room temperature. The membranes were then incubated with primary antibody overnight at 4°C in the buffer recommended by the antibody manufacturer. The membranes were then washed with TBS containing 0.2% Tween-20 (TBST) and incubated with species-appropriate fluorescently conjugated secondary antibody goat anti-mouse IRDye 680LT (1:10,000) or goat anti-rabbit IR Dye 800CW (1:10,000) from LI-COR Bioscience (Lincoln, NE, USA)] for 2 h at room temperature. Membranes were again washed in TBST and finally scanned to detect immunoreactivities using the Odyssey Infrared Imaging System (Li-Cor Bioscience). Data were quantified using pixel intensities with the Odyssey software (Li-Cor) according to the manufacturer’s protocols. The following antibodies were used at the indicated dilutions: Anti-OLIG2 (1:1000, MilliporeSigma, Burlington, MAcat# MABN50), Anti-MBP (1:500, MilliporeSigma, Burlington, MA; cat# SKB3-05-675), Anti-CASPR(1:2,000, MilliporeSigma, Burlington, MA; cat# MABN69), and Anti-MAG (1:1,000, Cell signaling, Danvers MA; cat# 9043S). Anti-β-ACTIN (1:20,000, Santa Cruz Biotechnology, Dallas, TX, USA; cat# sc-47778) was used as a loading control for all blots.

### Immunofluorescent staining

Mice were anesthetized with an *i.p*. injection of 750 mg/kg chloral hydrate and transcardially perfused with 4% paraformaldehyde in PBS pH 7.4. Their brains were post-fixed in PBS pH 7.4 overnight at 4°C, followed by PBS pH 7.4 with 30% sucrose for 72 h. 20 μm coronal brain sections were collected by cryosection for free-floating immunofluorescent staining. The sections were then incubated with the blocking solution (PBS pH 7.4 with 0.4% Triton X-100, 5% normal goat serum, 1% bovine serum albumin) for 2 h at room temperature, followed by incubation with the primary antibody. Primary antibodies: rabbit anti-OLIG2 antibody (1:1000, EMD MilliporeSigma, Burlington, MA; Cat# AB9610), mouse anti-OLIG2 antibody (1:1000, EMD MilliporeSigma, Cat# MABN50), mouse anti-CC1 antibody (1:500, Calbiochem, cat# OP80), mouse anti MBP Antibody (1:1000, EMD MilliporeSigma, Cat# 06-675), anti-GST-Pi antibody (1:1000, MBL international, Woburn, MA, cat# 312) or anti-PDGFRa antibody (1:1000, Cell signaling, Danvers MA, cat# 3461S). Sections were incubated with primary antibodies diluted in the blocking solution for 48 h at 4°C. Subsequently, the brain sections were washed in PBS 0.4% Triton three times and then incubated with goat anti-rabbit or goat anti-mouse Alexa Fluor-568, Alexa Fluor-488, or Alexa Fluor-647 secondary antibodies (1:800, Invitrogen, Waltham, MA) for 2 h at room temperature. 5-ethynyl-2’-deoxyuridine (EdU) was incubated using Click-iT™ Plus EdU Cell Proliferation Kit (Thermo Fisher Scientific) after DAPI staining. Sections were mounted with Prolong Diamond antifade mountant (Invitrogen, Waltham, MA). Three sections, representing rostral, medial, and caudal ACC (+.98mm, +.5mm, and −.10mm bregma), and hippocampus (−1.3mm, −1.8mm, and −2.5mm bregma) were analyzed for each set of staining. One image per hemisphere per bregma section for each animal was captured by a Leica TCS SP5 confocal microscope (Leica, Wetzlar, Germany) at 20x. Quantification was performed using the ImageJ software (U.S. National Institutes of Health) blinded to the experimental conditions using automated custom macro programs. For 5-ethynyl-2’-deoxyuridine (EdU) quantification, mice were injected intraperitoneally with EdU (80mg/kg) dissolved in 7.4 PH phosphate-buffered saline. To stain for EdU we used Click-iT™ Plus EdU Cell Proliferation Kit (Thermo Fisher Scientific) after DAPI staining on brain sections.

### Rat cannula implants and injections

Rats were anesthetized with ketamine (75 mg/kg) mixed with xylazine (10 mg/kg), and stainless-steel guide cannulas (C313G-SPC; 26-gauge P1 Technologies, Roanoke, VA) were implanted bilaterally using a stereotaxic apparatus (Kopf Instruments, Tujunga, CA) through holes drilled in the overlying skull to target the ACC (0.2 mm anterior, 0.5 mm lateral, −1.3 mm ventral from bregma). The guide cannulas were fixed to the skull with dental cement. Rats were administered meloxicam (3 mg/kg, subcutaneous) and let recover for at least 14 days before undergoing behavioral experiments. The injections were conducted using a 33-gauge needle, extending 1.5 mm beyond the tip of the guide cannulas, and connected via a polyethylene tubing (PE50) to a 1 μl Hamilton (Reno, NV) syringe controlled by an infusion pump (Harvard Apparatus. Holliston, MA) 2 nmol of antisense oligodeoxynucleotides (AS-ODN) or the relative scrambled sequence (SCR-ODN) were delivered per brain hemisphere in 0.5 μl of PBS (pH 7.4) at a rate of 0.333 μl/min. Sequences were as follows: Myrf AS 5′-GGTCTCGTCCACCACCTCCAT-3′; Myrf SCR 5′-CCATCTTCCGACGTTCGACCC-3′. The SCR-ODN, which served as control, contained the same relative AS-ODN base composition but in random order and showed no homology to any mammalian sequence in the GenBank database, as confirmed using a basic local alignment search tool (BLAST). All ODNs were phosphorothioated on the three-terminal bases at each end to protect against nuclease degradation. ODNs were synthesized, reverse-phase cartridges purified, and purchased from Gene Link (Hawthorne, NY). Rats were euthanized at the end of the behavioral experiments to confirm cannula and injection placement. Toward this end, 40 μm coronal sections were sliced following fixation of the brains in 10% formalin; then, the sections were examined under a light microscope to verify cannula placement. Rats with incorrect placement were excluded from the study.

### Object location memory

Mice were habituated, trained, and tested in a square, open field (29 × 29 × 18 cm) with white Plexiglas walls and floor measured at 12.5 (±2.5) lux in the center of a dim room. Visual cues were provided within the box and on the walls of the room. Behavior was recorded with a video camera positioned above the arena. Mice were first habituated to the arena for 10 minutes for 3 consecutive days before the training. Twenty-four hours after the last habituation session, each animal was returned to the arena for its training session. Training consisted of exposing the mice to two identical objects constructed from Mega Bloks (Montreal, Canada) secured to the floor of the arena. Object sizes were no taller than twice the size of the mice. Mice were initially placed facing a corner, away from the objects, and were allowed to explore the arena and objects for 10 min. 4 hours after training; each animal was tested in the arena. During testing, one object remained in the same location as during training, whereas the second object had been moved to a novel location. Animals were placed in the arena facing the same direction as during training and were allowed to explore for 10 min. The placement of the object in the novel location was counterbalanced between subjects. The arena and objects were cleaned between sessions. Video files were coded and scrambled. The experimenter was blind to treatment and scored the total time the mice spent actively exploring each object in each session. Active exploration was defined as the mice pawing at, sniffing, or whisking with their snout directed at the object from a distance of less than ~1 cm. Sitting on or next to an object was not counted as active exploration. Mice with less than 10s total exploration time were excluded. If mice explored more than 15s, the exploration percentage was taken at 15s of total exploration time. Memory was measured as the percentage of time spent exploring the object in the novel location compared with the stationary object.

### Open field

Mice were allowed to freely explore an open-field arena illuminated at 195 lux. (43.2 cm × 43.2 cm × 30.5 cm (Med Associates Inc., St. Albans, VTENV-515) for 10 min. The open field was designated into 2 sections: center box and outer border. Percentage time spent in the center and average velocity and total distance were quantified. Activity was analyzed with Ethovision-XT (Noldus Information Technology).

### Mouse viral injections and C21 administration

Mice were anesthetized with isoflurane. The skull was exposed, and holes were drilled in the skull bilaterally above the ACC or dHC. A Hamilton (Reno, NV) syringe with a 33 gauge needle, mounted onto a nanopump (K.D. Scientific, Holliston, MA), 0.2ul microliters of the virus was injected per mouse bilaterally into the ACC (+ 0.5mm anterior to bregma, ± 0.3 lateral of bregma, −2 dorsal of skull surface) or 1ul per mouse bilaterally into the dorsal hippocampus (+1.7mm anterior to bregma ±1.5 lateral of bregma −1.75 dorsal of skull surface) at a rate of 0.2μL/min. The injection needle was left in place for 5 min following injection to allow complete dispersion of the solution and then the scalp was sutured. Meloxicam (3 mg/kg) was used as an analgesic treatment after surgeries, and mice were allowed to recover for 14 days before training.

The pAAV-*Mbp*-CreER^T2^ virus (titer: 10×13 GC/μl) was packaged into AAV-PHP.B capsid and purchased from Vector Biolabs (Malvern, PA, cat# VB1545). The AAV-hSyn-hM4D(Gi)-mCherry (cat# VB1545) was purchased from add gene (titer: 7×10^12^ vg/mL; cat# 50475-AAV8). C21 (HB6124, Hello Bio, Princeton, NJ) was dissolved in PBS pH7.4 and injected at 1mg/kg 60 min before training. After behavioral experiments, mice were anesthetized with an *i.p*. injection of 750 mg/kg chloral hydrate and transcardially perfused with 4% paraformaldehyde in PBS pH 7.4. Their brains were post-fixed in this solution overnight at 4°C, followed by PBS pH7.4 with 30% sucrose for 72 h. 30 μm brain sections were collected by cryosection for free-floating immunofluorescent staining.

### Statistical analyses

Data were statistically analyzed using Prism software. The student’s t-test was used to compare statistical differences between two experimental groups. When more than two groups were compared, data were analyzed with one- or two-way repeated-measure ANOVA followed by Bonferroni post hoc test. All values represent the mean ± standard error of the mean (SEM). The experimental n, the statistical test used, and the statistical significance are indicated in figure legends. The Excel-based PCR Array Data Analysis was used to analyze the qPCR results. The number of independent experiments carried out and the numbers of biological replicates [i.e., animals (n)] are indicated in each figure legend. No statistical method was used to predetermine sample size. The numbers of subjects used in our experiments were the minimum required to obtain statistical significance, based on our experience with the behavioral paradigm and in agreement with standard literature.

## Acknowledgments

We thank Dr. James Salzer (New York University School of Medicine) for providing an initial group on transgenic mice and for helpful discussions. This work was supported by NIH grants R37MH065635 to CMA, HHMI Gilliam fellowship to LPB. ON was supported by NIGMS MARC grant 5T34GM008078.

## Author contributions

LPB, CMA, designed the study. LPB, BB, and ON performed the experiments. LPB and CMA wrote the manuscript.

## Ethics

All animal procedures complied with the US National Institute of Health Guide for the Care and Use of Laboratory Animals and were approved by the New York University Animal Care Committees. All surgeries were performed under isoflurane anesthesia and every effort was made to minimize suffering.

---

**Supplementary data Figure 1.**
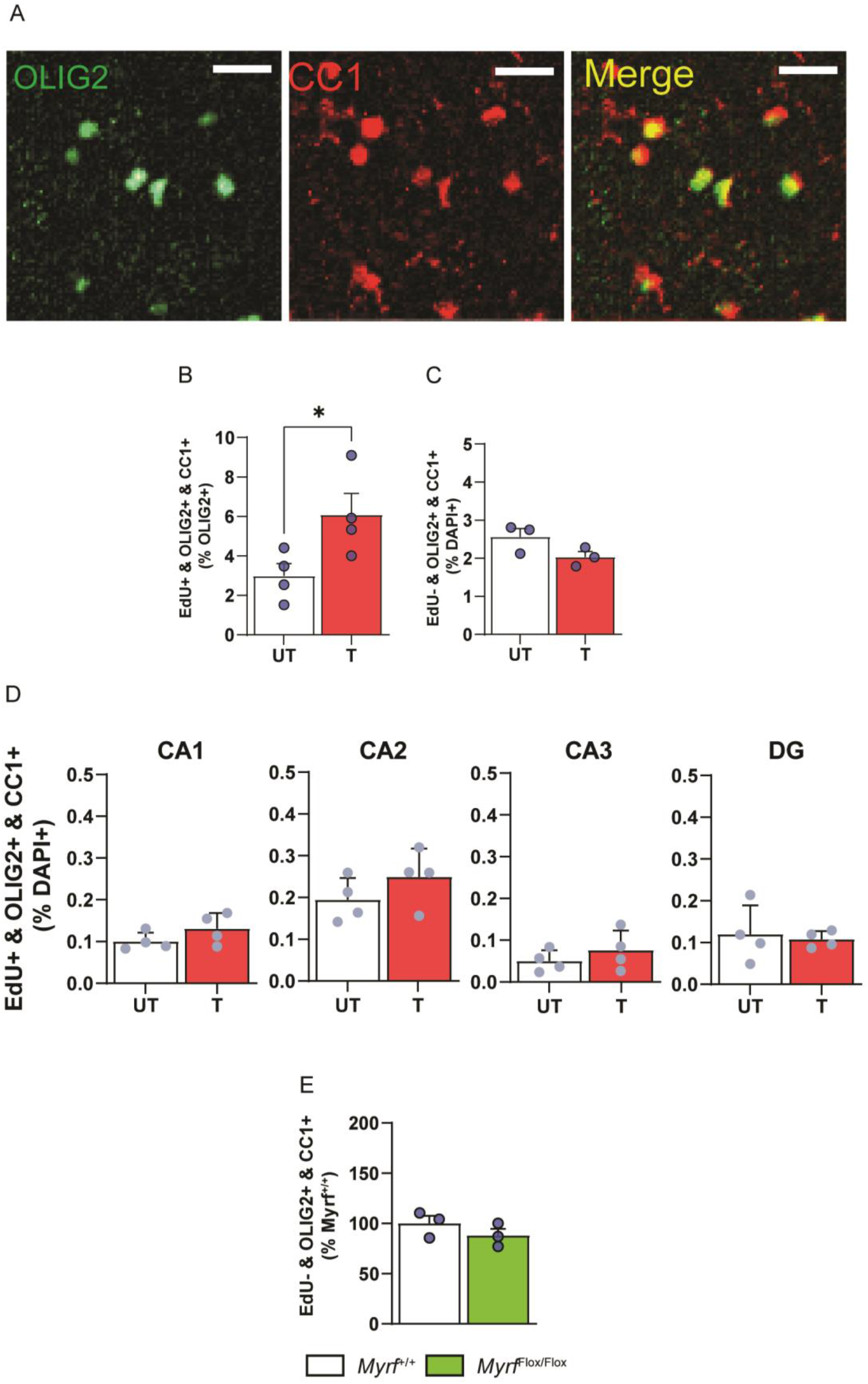
(A) Representative images are taken from Fig. 4A: immunofluorescent staining of OLIG2 and CC1 in the ACC of rats euthanized 1 day after IA compared to UT control. Here OLIG2 intensity was increased to show detection of colocalization with CC1 (scale bar:40 μm). (B) Quantifications of ACC triple immunostaining of EdU, OLIG2 and CC1 relative to OLIG2+ cells (n = 4 mice per group, two-tailed t-test; * Indicates p<0.05). (C) Quantifications of ACC triple immunostaining of EdU−, OLIG2+ and CC1+ relative to DAPI+ cells (n = 3 mice per group, two-tailed t-test). (D) Quantifications of EdU, OLIG2 and CC1 triple staining in the dHC subregions CA1, CA2, CA3, and DG relative to DAPI. Three coronal sections were quantified and averaged. In each coronal section the entire CA1, CA2, CA3, or DG were quantified bilaterally. Each dot in the graphs represents the average values of the three coronal sections of each mouse. B-E: Data are presented as mean percentage ± s.e.m. of positive cell number relative to OLIG2+ cells or DAPI+ nuclei as indicated in each graph (n = 4 mice per group, two-tailed t-test). For detailed statistical information, see table 2-source data1. (E) Quantifications of ACC triple immunostaining of EdU-, OLIG2+ and CC1+ relative to DAPI+ cells (n = 3 mice per group, two-tailed t-test).

